# Unraveling the network signatures of oncogenicity in virus-human protein-protein interactions

**DOI:** 10.1101/2025.01.14.632978

**Authors:** Francesco Zambelli, Vera Pancaldi, Manlio De Domenico

## Abstract

Climate change, increasing urbanization and global human mobility have increased the risk of emergent infectious diseases and the risk for their pandemic potential. Consequently, there is a pressing need for methods that can provide rapid insights about the potential enduring effects on the human hosts. *In-silico* methods can be more suitable than *in-vitro* and *in-vivo* methods to identify effects potentially manifesting after a long time, under the hypothesis that the underlying causes can be ascribed to perturbed biomolecular processes. Here we focus on oncogenicity, i.e., the risk of developing cancer subsequent to a viral infection, a phenomenon estimated to account for approximately 15% of global cancer prevalence. We characterize viruses in terms of their oncogenic potential by analyzing a multilayer representation of protein-protein interaction networks reconstructed from the human interactome. Our framework facilitates classification and allows us to identify sub-sets of proteins in-volved in oncogenesis, highlighting the mechanisms underpinning viral oncogenicity, including factors like chromatin structure. Our framework serves as a foundational resource for medical research, paving the way for the study of other potential long-term effects associated with emerging viruses.

## Introduction

In the last years, the profound impact of viral pathogens on human health has become unmistakably evident, with the ongoing global pandemic serving as a stark reminder of their relevance. Beyond the acute phase of an infection, it is increasingly recognized that certain viral pathogens can cause persistent health issues[1, 2, 3, 4], notably among them, the potential for oncogenicity [5, 6, 7, 8, 9], which poses a substantial and multifaceted challenge for both individuals and public health. Such a risk is further emphasized by recent discoveries about cellular mechanisms that lead to DNA degradation and cell senescence subsequent to SARS-CoV-2 infection[10].

In the last two decades, the study of complex networks has undergone significant advancements, which have broadened its impact on various fields. One of the major breakthroughs concerns the application of complex network theory to systems biology and medicine, leading to network medicine [11], an interdisciplinary field that enables studying diseases and their underlying mechanisms in terms of network structure and dynamics[12, 11, 13, 14]. This approach focuses on understanding phenotype in terms of the interdependence between proteins, genes, metabolites, biological processes and drugs, providing a suitable framework to build *in silico* models, shedding light on the underlying biological systems and on complex disease-disease interactions [15, 13, 16].

One of the core elements used in network medicine are protein-protein interaction (PPI) networks[17, 18, 19, 20, 21], models able to represent the interactions between proteins in a biological system. These interactions can be physical (direct) or functional (indirect) and are crucial for a variety cellular processes and functions[22, 23]. PPI networks provide a way to visualize and study how proteins interact with each other, forming complex relationships within a cell or organism.

Recent work exploited such tools to gain deeper knowledge about the Sars-Cov2 virus [24, 25, 26, 27], for example allowing integration of interaction between proteins, symptoms and biological pathways to devise drug repurposing strategies[28, 18].

Considering the increasing number of results obtained from the study of complex diseases by means of PPI networks, we ask whether it is possible to apply such methods to determine the potential oncogenicity of a virus following virus-host interactions. The wide availability of this type of PPI data[22] offers a a valid starting point to tackle this challenge.

Moreover, the prospect of utilizing online data and mathematical methodologies to establish a classification system for viruses based on their oncogenic potential holds significant promise for systems medicine. In fact, the outcome of transparent computational models, based on network analysis, can not only serve as an invaluable springboard for ongoing clinical and medical investigations, but also provide valuable insights into the effects of emergent viruses, which have yet to undergo comprehensive scrutiny due to their relatively short existence and the lack of longitudinal data. A salient case in point is the SARS-CoV-2 virus, for which evidence about possible long term effects on human organs and systems are already under investigation [29, 30, 31]. In this case, a complete examination of the whole spectrum of its long-term effects from a clinical perspective might require at least a decade.

To address this challenge, we use a multilayer framework [32, 33] where multiple virus-host PPI networks are coupled to build multiplex networks [34, 35]. Systems consisting of multiple interacting networks have attracted considerable attention, owing to the discovery of novel structural and dynamical features in coupled cases that differ from those observed in uncoupled representations of the same system, as reported for transportation [36], urban [37], ecological [38], social [39], financial [40] and many other systems [41, 34, 42, 43, 44], to mention some emblematic examples (see [33] for a recent review).

Here, we use multilayer network models to construct a holistic system able to account for perturbations due to distinct external agents (i.e., viruses) to the same target (i.e., human cells). Considering the whole human interactome as a proxy for this target, each layer can be understood as a description of the interaction with a specific virus.

In fact, this framework facilitates the identification of commonalities and shared properties between these distinct layers of interaction with the human proteins, being suitable to providing insights into shared characteristics, such as oncogenicity.

To this aim, we develop a two approaches to address a classification task. We start by computing a diverse set of multilayer topological features, focusing on the ones that allow us to statistically distinguish networks sampled from ensembles of oncogenic and non-oncogenic viruses. Then machine learning techniques were exploited to perform a classification between virus-host interaction PPI networks associated to oncogenic and non-oncogenic viruses, and by doing so, they allow also to identify sets of proteins that play a significant role in predicting oncogenic potential.

By applying functional enrichment analysis to this set of pivotal proteins, a noteworthy emphasis emerges on pathways intimately linked to chromatin structure, a phenomenon extensively associated with the onset of cancer as evidenced by numerous studies[45, 46, 47]. Additionally, we observe a convergence with well-established cancer-related pathways, including the WNT pathway, TP53, and pathways encompassing sumoylation processes, all of which play pivotal roles in our quest to comprehend viral oncogenic mechanisms.

## Results

The work is based on the analysis of PPI data sourced from online databases and processed to create models describing the interaction between viruses and host. Successively multiple PPI networks corresponding to different viruses, are combined into a multilayer network framework, creating the expanded dataset to be used in the analysis phase.

### PPI networks as proxy for virus-human interactions

This study relies on the BIOstring database[48], a large repository of protein-protein interaction (PPI) data that has been curated by integrating information from various sources, including BioGRID [49] and STRING[50, 51]. Specifically, the database contains PPI interactions between 80 viruses and human cells, where 8 viruses are categorized as oncogenic and 72 as non-oncogenic based on clinical data. Note that this data represents a composite, “mean field” model, as the PPIs are collected under diverse conditions and may involve various human cell types.

Concerning virus-host interactions, the core concept is that when a virus protein targets a human protein, the resulting effects can vary[52, 53]. These effects may include inhibiting or disrupting certain interactions while creating new ones. Instead of predicting how the human interactome might change, the focus here is on identifying the region of this large network that is most likely to be affected. By studying this specific subset of the whole human interactome, we aim to gain insights into the strategies employed by the virus to attack a human cell.

The construction of the PPI networks associated to each virus is described in the top section of Figure 1 and further details are provided in the Material and Methods section. At the end of such procedure, we end up having a set of 80 PPI networks, each associated to a specific virus, describing their interactions with the human cell.

**Figure 1:**
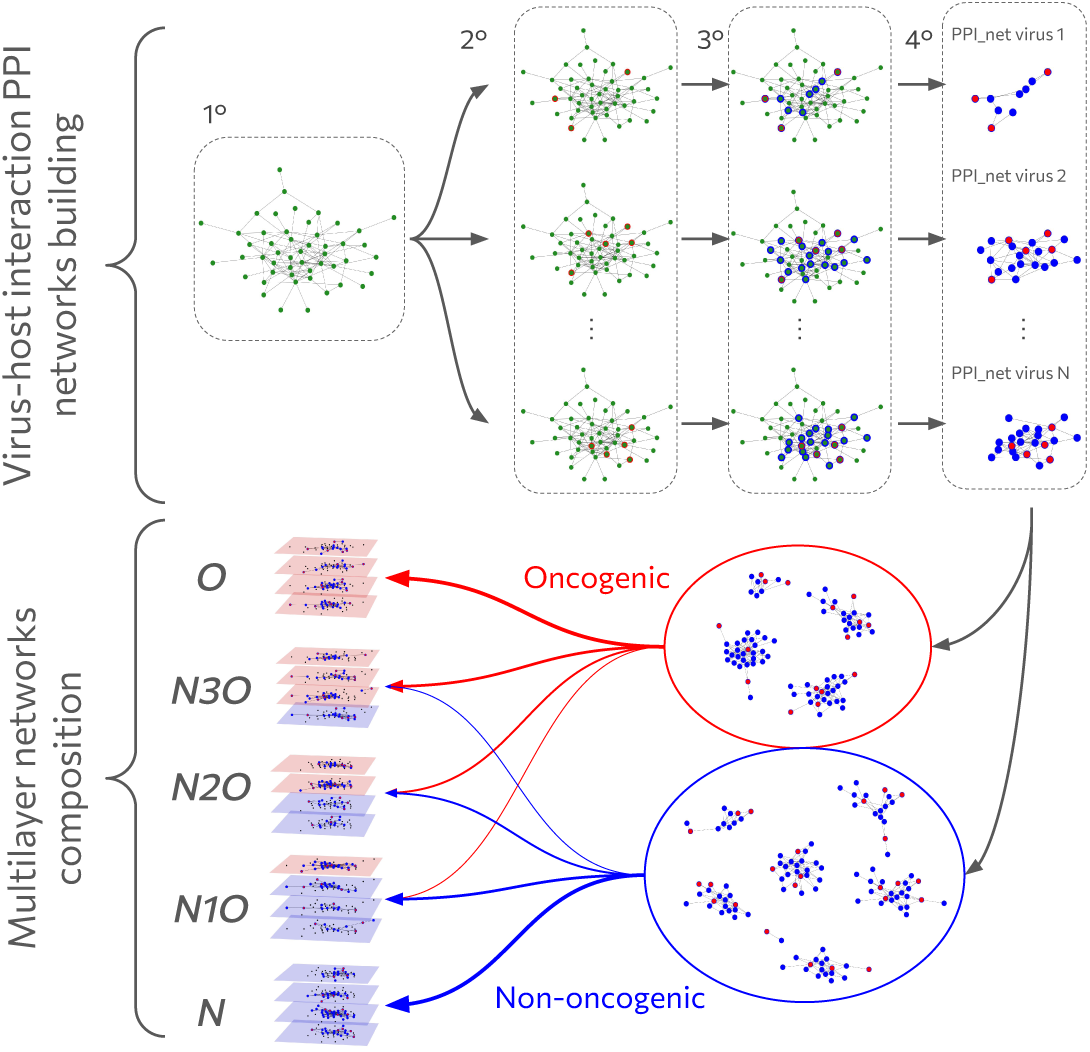
Schematic illustration of the workflow. The figure represents the two main phases of the creation of the analysis dataset. In the first one, depicted in the top of the figure, an illustrative example demonstrates how the Protein-Protein Interaction (PPI) networks describing the interaction between each virus and the human interactome are built. The first step consists of considering the entire human PPI, built by integrating PPI data from different data sources. A visual example is provided by the figure in the top right of the plot, showing a network with green nodes. In the second step, for each virus, the human proteins directly targeted are selected, and they correspond to the nodes highlighted in red in the figure. To expand the number of nodes likely to be influenced by the virus, in the third step, the nearest neighbors of the directly targeted proteins are added, which are highlighted in blue in the figure. Finally, in the fourth step, the human interactome is subset only to the highlighted nodes, resulting in a set of PPI networks, each representing the interaction network between a virus and the host. The second phase consists of building multilayer network samples. In particular, each of the PPI networks created earlier can be classified as being associated with an oncogenic or non-oncogenic virus, based on the a priori information retrieved in the data collection phase. These networks are then combined in groups of four, considering all possible combinations of oncogenic and non-oncogenic networks to be grouped. Each group of networks is then organized in a multiplex framework, creating a dataset to be used in further analyses.

### Combining multiple virus-host interaction PPI networks to uncover features shared by several viruses

To exploit the full potential of PPI networks, we employed a multilayer framework, which involves combining a fixed number of virus-host interaction PPI networks over the total 80 belonging to our dataset, in a layered structure, in which each layer corresponds to one of the chosen networks. This approach does not discard, *a priori*, any available information, and it has been used in a variety of disciplines to show that it is often better than considering layers in isolation or aggregating them (see [33] for a recent review).

The multilayer framework serves several key objectives. Firstly, it allows us to interpret each layer as a distinct source of external perturbations – specifically, distinct viral infections – on the human interactome. Secondly, by combining multiple systems (i.e., virus-host PPI networks) that share a common feature, we can identify common properties among these individual systems, which underlie this common feature of interest. Lastly, this framework allows us to generate a wide array of unique samples, built by combining fixed-size sets of virus-host PPI networks. Having generated a high number of these sets, it is possible to ‘robustly’ investigate the statistical distributions of various descriptors extracted from these networks.

Such a procedure is represented graphically in the bottom panel of Figure 1, while mathematical details are provided in the Material and Methods section.

We chose to include 4 virus-host PPI networks in each multilayer network sample, representing a trade-off between the computationally complexity of the resulting object (each virus-host PPI has a number of nodes of order 10^4^ and 7 · 10^5^ edges), and the possibility generate a large number of samples, resulting in combination of virus-host interaction PPI networks associated to different viruses. In fact, the larger is the size of such groups, the larger is the number of possible combinations. Multilayers with 4 layers appeared to be the best trade-off between these 2 aspects.

As reported before, each virus is classified as belonging to the oncogenic or non-oncogenic class, thus each 4-element combination of virus PPI networks is grouped based on the number of oncogenic and non-oncogenic viruses in it.

In the following, the various combinations of oncogenic and non-oncogenic layers used to create these samples are referred to as “combination sets”. Each combination set is denoted by a label that indicates the number of layers of each type it includes. For simplicity, we also designate layers associated with virus-host interaction PPI networks linked to oncogenic viruses as “oncogenic layers”, and similarly for “non-oncogenic layers”.

Our aim is to study (dis)similarities between descriptive quantities that can be extracted from the different combination sets, thus revealing if the different proportion of oncogenic and non-oncogenic layers can be a statistically discriminating feature over the data. Table 1 reports combination sets considered here, detailing the number of oncogenic and non-oncogenic layers, as well as the total number of possible samples that can be generated from each class, and the corresponding abbreviations used for reference.

**Table 1:**
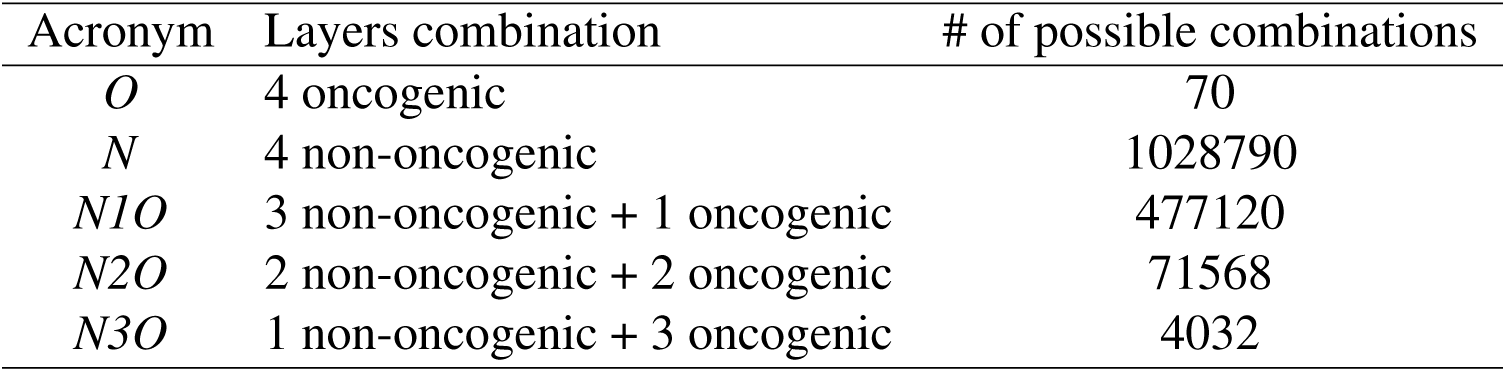
Combination sets used in this study, built from combining oncogenic and non-oncogenic layers. The number of possible different samples that can be obtained from the data is reported in the last column.

| Acronym | Layers combination | # of possible combinations |
| --- | --- | --- |
| *O* | 4 oncogenic | 70 |
| *N* | 4 non-oncogenic | 1028790 |
| *N1O* | 3 non-oncogenic + 1 oncogenic | 477120 |
| *N2O* | 2 non-oncogenic + 2 oncogenic | 71568 |
| *N3O* | 1 non-oncogenic + 3 oncogenic | 4032 |

### Network features provide biological insights about virus-host PPI systems

We build ensembles of samples from different combination sets and, for each one, we calculate a variety of topological features potentially associated to some underlying biological features of the corresponding systems. Therefore, we compare the statistical distributions resulting from different combination sets and search for statistically significant differences.

### Identifying the core of nodes biologically relevant for oncogenesis

First, we set out to define a framework to identify core nodes in the multilayer networks, which normally constitute a component. In multilayer networks, the concept of what a component is depends on which specific features of the system should be considered. Importantly, all these definitions represent core structures within the system, making them highly relevant[54]. We studied 3 types of components: the largest connected component (LCC), corresponding to the maximum subset of nodes connected by at least one multilayer path between each other, the largest intersected component (LIC), in which nodes must be connected in all the layers independently, and the largest viable component (LVC), where the set of nodes must be connected by the same path in all the layers independently. In particular, nodes belonging to the LVC are usually considered to be very important, because they are crossed by paths which are present simultaneously in all the nodes, and for this reason it is very likely that they are fundamental to determine the overall system behaviour[55, 56, 57].

By studying how these cores change across combination sets, we gain quantitative insights about how different viruses interact with the same regions of the human interactome. Here, we focus on investigating the size of such components, and statistically comparing them to draw conclusions.

The first analysis includes the study of the LCC sizes. We found that they are essentially proportional to the network size, thus not providing a relevant feature for classification, which tends to differentiate only between large and small virus-host interaction networks rather than focusing on more distinctive properties (see the Material and Methods for details).

Conversely, the LIC and LVC offer more valuable insights. They uncover common structures across different layers, i.e. different virus-host interaction patterns, enabling the identification of shared properties across layers. The focus in our case falls in the oncogenic-non-oncogenic composition of the combination set.

Firstly, we studied the distribution of their size, which quantifies how many target nodes are shared by the viruses whose associated PPI networks compose the multilayer network. The statistical distributions of the LIC and LVC sizes are highly similar, often differing by just a few nodes, and thus in the subsequent analysis, only the LVC case is considered. Figure 2 a) shows the box plot representing the distributions of the LVC size for each combination set. Notably, in the oncogenic case the distribution of these values skews toward higher values, indicating that oncogenic viruses tend to influence similar and overlapping regions of the human interactome. A Wilcoxon rank-sum test, comparing the alternative hypothesis of the *O* distribution being stochastically greater than the *N* one, yields a p-value of 5 · 10^−15^, revealing a strong statistical difference between the two extreme cases.

**Figure 2:**
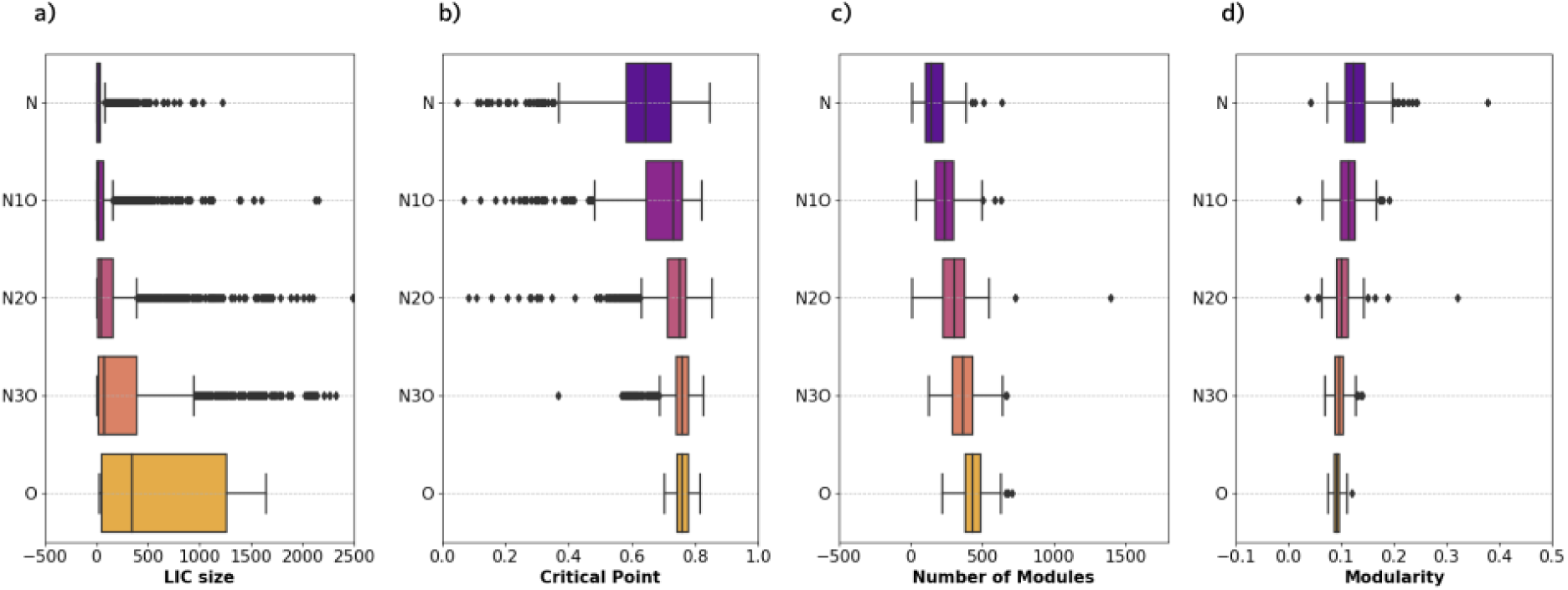
Statistical distribution of the topological features extracted from the multilayer network samples: The figure displays box plots illustrating the distributions of select topological features extracted from the various multilayer network samples within each combination set. For each of those, the colored box represents the first quartile of the distribution, while the line traversing it is the median. The The “whiskers” extend to points that lie within 1.5 IQRs of the lower and upper quartile, and then observations that fall outside this range are displayed independently. Specifically, in (a), we visualize the distribution of the size of the largest intersected component. Moving on to (b), we observe the critical point, denoting the fraction of removed nodes at which the transition phase occurs during node percolation. This transition is determined by following the order indicated by the multipage rank versatility computed for each network. In (c), the number of communities or modules is showcased, while in (d), the figure presents the modularity value retrieved with the DCSBM community partition algorithm.

Another crucial finding is that when comparing the *O* and *N3O*, the same type of rank-sum test yields a p-value of 2 · 10^−3^, indicating that even inserting just a single non-oncogenic layer into a multilayer network primarily composed of oncogenic virus layers results in a clearly distinguishable distribution from the case of 4 oncogenic layers.

To gain deeper insights into the characteristics of oncogenic viruses, we investigate a common core of nodes shared by all the PPI networks associated with them. We accomplish this by extracting the node sets corresponding to the LIC from each sample generated from the *O* combination set and identifying the common nodes. This process corresponds to finding the LIC within an 8-layer multilayer network constructed from all the PPI networks associated with oncogenic viruses. The outcome is a list of 30 proteins -extensively reported in S8 Appendix-, highly enriched for pathways involved in transcription, DNA-binding and sumoylation processes.

### Robustness of the human interactome to viral targeted attacks

Another significant feature that can extracted from each network is its critical point in response to a targeted percolation process (see [58] for a recent review). In the context of network science, percolation can be interpreted as a way to simulate an attack process aimed at network dismantling. Here, we use the multilayer pagerank versatility [59, 60] to rank the nodes to be used for targeted attacks (see Materials and Methods for mathematical details). From a biological perspective, this quantity identifies the proteins that attract the majority of information flows (under the assumption of a diffusive process), indicating potential functional relevance.

The critical point corresponds to the minimum fraction of nodes to be attacked to disintegrate the largest connected component of the system. Therefore, it provides insights into the network’s responsiveness to external disturbances and, in the context of this study, it is related to how vulnerable the virus-influenced region of the interactome is to external perturbations. Observing the size of the largest and second-largest components of the network as nodes are removed allows us to identify the fraction of removed nodes at which a phase transition occurs. This transition marks the shift from an ordered phase, where a dominant, well-connected component governs the system, to a disordered phase, where no single dominant cluster of connected nodes exists.

Analyzing the distributions of the node percolation critical points from various combination sets, reported in Figure 2 b) reveals a noteworthy trend: a shift towards higher critical point values as the number of oncogenic layers in the combination set increases. Specifically, a Wilcoxon rank-sum test considering the alternative hypothesis of the *O* distribution to be stochastically larger than the *N* distribution, yields a p-value of 9 · 10^−13^, which allows us to reject the null hypothesis, and therefore shows clearly that interactome subsets impacted by oncogenic viruses are more resilient to this type of targeted attack.

### Modular structure is related to oncogenicity

The final set of features under analysis are those extracted from community detection [61, 62], a method focused on identifying groups of nodes that are organized at the mesoscale, according to some prescription, from modularity maximization[63, 64] to inferring the parameters of a generative model such as the degree-corrected stochastic block model (DCSBM) by means of a Bayesian approach [65]. In this work we use the second method, that has been also reliably extended to multiplex networks like the ones we have built [66]. In our study, this analysis offers insights into the network’s dispersion: the greater the number of communities within the system, the more likely it is that a higher number of functions are associated with the set of nodes in the network, indicating higher functional differentiation and consequently a more prominent systemic behaviour. Conversely, a smaller number of communities might suggest that most nodes are linked to similar functions. Of course, this is true under the hypothesis that groups of proteins corresponds to modules with special structural or functional meaning, e.g., specific biological processes within a cell. This is true is we further assume that these interactome subsets we are considering are similar to the original full interactome, in which network communities correspond to biological processes or genes involved in the same functions[67, 11].

Another quantity of interest is modularity [68]: the higher the modularity, the more segregated the communities are, suggesting fewer connections between them and a more marked group mesoscale structure.

The results shown in Figure 2 c) and d), clearly indicate an inverse relationship between the number of modules and modularity when considering changes in the number of oncogenic layers within the combination set. As the number of oncogenic layers increases, the number of modules significantly rises, while the modularity distributions shift towards lower values. This observable trend can be quantitatively assessed using Wilcoxon rank-sum tests.

In particular, when we test the alternative hypothesis that the *O* distribution has a larger number of modules compared to the *N* distribution, the test yields a p-value of 2 · 10^−13^, providing strong evidence against the null hypothesis. Similarly, when we consider the modularity, the test is performed with the opposite hypothesis, evaluating whether the *O* distribution is smaller than the *N* distribution. This test also gives a p-value of 2 · 10^−13^, allowing us to confidently reject the null hypothesis.

Overall the number of modules appears to be statistically greater for the *O* distribution compared to the *N* one, while the opposite happens for modularity, in which the distribution associated to *O* results to be statistically smaller than the *N* one.

### Exploration of combination sets based on dimensionality reduction of topological parameters

After analyzing each of the considered features separately, as described in the previous paragraphs, the objective is to combine these different quantities to check if even with this method it is possible to propose a strategy to distinguish samples belonging to the different combination sets. To accomplish this, the UMAP algorithm, a dimensionality reduction technique centered on unsupervised clustering, is employed. UMAP aims to maintain the global structure of high-dimensional data while reducing it to a lower-dimensional space. It was chosen between the other analogous algorithms for its ability to detect non-linear relations. Out of the six extracted features, only four were utilized in this phase. As demonstrated earlier, LCC dimension is closely tied to network size and could introduce bias, while the dimensions of LIC and LVC are highly similar, offering redundant information. The final set of features used to differentiate the parameter space includes percolation critical point, LVC dimension, the number of non-empty communities, and modularity for each multilayer network.

Using UMAP, each point is projected onto a 2-dimensional space. Subsequently, a support vector machine algorithm with a Gaussian kernel is applied to separate the reduced parameter space into two regions: an “oncogenic region”, associated to samples containing only oncogenic layers, and the “non-oncogenic region” associated to samples composed only by non-oncogenic layers.

The results are displayed in the top row of Figure 3, and they show a clear distinction between the samples belonging to the two classes. Remarkably, no fine-tuning of the dimensionality reduction algorithms was necessary to achieve such result.

**Figure 3:**
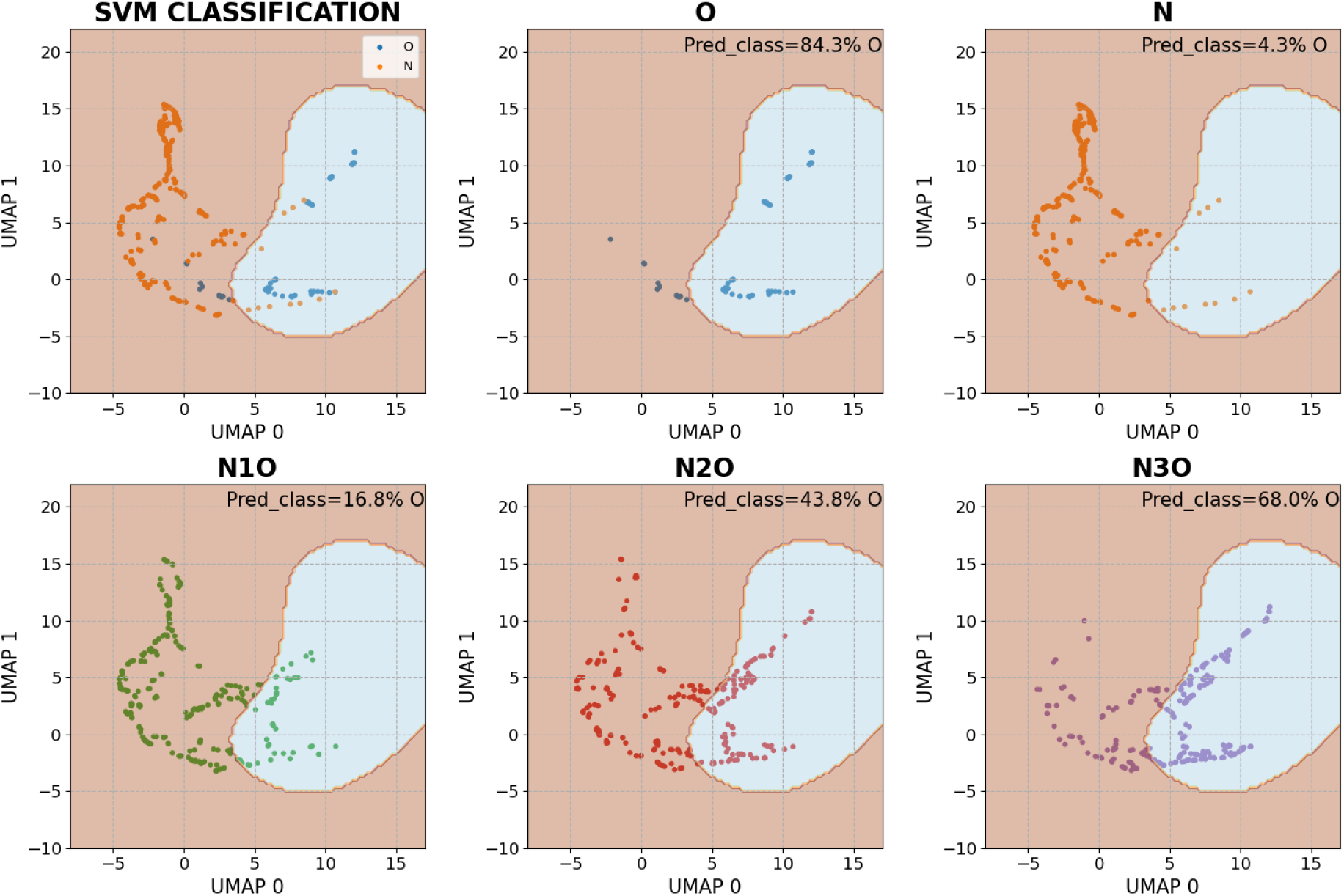
Combined feature classification: Each point in this plot represents a 2-dimensional projection generated using UMAP derived from the 4-dimensional vectors composed of various topological features linked to multilayer networks within one of the combination sets. These features include LIC size, percolation critical point, number of modules, and modularity. In the top-right plot, points corresponding to samples from category *O* (in blue) and *N* (in orange) are presented. The parameter space is effectively divided into two distinct regions, one associated with samples from category *O* (shaded in blue) and the other with category *N* (shaded in orange). This classification is achieved by employing a Support Vector Machine (SVM) with Gaussian kernels. The remaining five plots illustrate how the points generated from different combination sets are distributed within the parameter space. For each of these plots, the fraction of points belonging to the region associated with samples from category *O* is reported. It becomes evident that this fraction decreases as the number of non-oncogenic layers associated with the combination set increases.

After the separation of the reduced parameter space in the two regions, we also consider samples generated from the remaining combinations. The 4-features vectors associated to each of their samples is projected into the 2D space using the same UMAP algorithm, falling either in the “oncogenic” or “non-oncogenic” region. In Figure3 we graphically represent the distribution of the points associated to the samples from each combination set alongside to the fraction of such points falling in the “oncogenic” region. The expectation is that, as the fraction of layers associated with oncogenic viruses increases, the dimensionally reduced points will shift from the “non-oncogenic” to the “oncogenic” region. This expectation is validated by the results depicted in the three images in the bottom row of Figure 3. For each combination set, the fraction of samples classified as belonging to the region associated with the *O* combination set is also reported, and this value increases as the number of oncogenic layers grows significantly.

The fact that the fraction of samples classified as belonging to the *O* region exhibits significant change with the ones from *N1O*, suggest that the algorithm is sensitive to the introduction of a single layer of the non-oncogenic type. It is reasonable to expect that such clear transitions would persist when introducing a new virus, enabling classification through comparison with the known scenario, even in the case of entirely novel viruses.

### Functional validation of findings

At this point the following question could arise: are these results coming from a model that actually contains information about the human cell and its interaction with viruses, or just to the connectivity of human proteins in the interactome?

To investigate this further, we compared the results obtained above with results obtained after a randomization procedure applying random rewiring using the configurational model of the complete human interactome, before extracting the virus-host interaction networks. From this we apply analogous analyses to the ones described in the previous section and compare them statistically, looking for significant differences. Such procedure is repeated multiple times to create a representative null model. Further details about the procedure are reported in Material and Methods.

What we find is that LVCs from randomized models are statistically much less functionally enriched with respect to the one of the original dataset -see S3 Appendix-. The percolation analysis reveals that for the randomized model the networks are statistically less robust -see S4 Appendix-, while the values of the modularity suggests that rewiring the original human interactome, the resulting multilayers composed of virus-host interaction networks are characterized by a weak block structure -see S5 Appendix-. All these findings strengthen the claim that the dataset over which the analysis is performed indeed contains biological relevant information about the relationships between viral proteins and specific human cell processes, and thus the previous and following analysis can be considered meaningful.

### Classification of oncogenic vs non-oncogenic virus-host interaction PPI networks using machine learning

After proposing a method to distinguish virus-host interaction multilayer networks composed by different number of oncogenic and non-oncogenic layers based on topological features, we aim to explore methods to perform classification of individual viruses. This approach seeks to determine whether a single virus-host interaction PPI network is associated with an oncogenic virus or not. An important secondary objective is to identify key proteins relevant for classification, potentially shedding light on mechanisms underlying virus oncogenicity.

To accomplish this we exploited the power of machine learning classification algorithms, using as inputs the vectors produced from an embedding procedure of the multilayer networks used in the previous section. Such an embedding assigns to each node a value proportional to its multi page-rank versatility, creating a different fixed size vector for each possible sample in the dataset. The pipeline is described in more detail in the Supplementary material section.

Using the dataset from the previous analysis sections composed of 4-layer multilayer network samples, our chosen strategy to identify the contribution of a single layer associated with a specific virus focuses on classifying samples composed only by oncogenic layers, from the *O* combination set, from ones containing only one non-oncogenic layer, thus belonging to *N1O*. This choice follows the fact that by considering such combinations sets, it is possible to have training and test sets large enough to perform a training procedure. If the performance of the classification algorithms is accurate and exhibits good generalization capabilities, it should also possible to propose a classification for PPI networks associated with viruses not used for training by considering them alongside the oncogenic viruses used for the training.

Alongside classification performance, another important requirement for the machine learning algorithm is interpretability. In our case, this means being able to extract the most relevant input features for the classification. This allows us to achieve our secondary goal, which is to extract a set of proteins relevant for distinguishing between oncogenic and non-oncogenic cases. In this context, we chose a perceptron model, as it is a powerful yet easy-to-interpret machine learning algorithm.

To assess the algorithm’s generalization capabilities and ensure results are not influenced by overfitting, the perceptron is trained and tested multiple times across different datasets, producing different final algorithms. Nevertheless, they all share the same classification objective. In each trial, the training set retains samples from the *N1O* combination set, which contains an oncogenic layer associated with a specific oncogenic virus. Subsequently, the algorithm’s performance is evaluated on this sample set to determine if it can recognize an oncogenic virus, even if it was not part of the training data.

As reported in Table 2, the performances are strong in nearly all trials, resulting in very high classification precision.

**Table 2:**
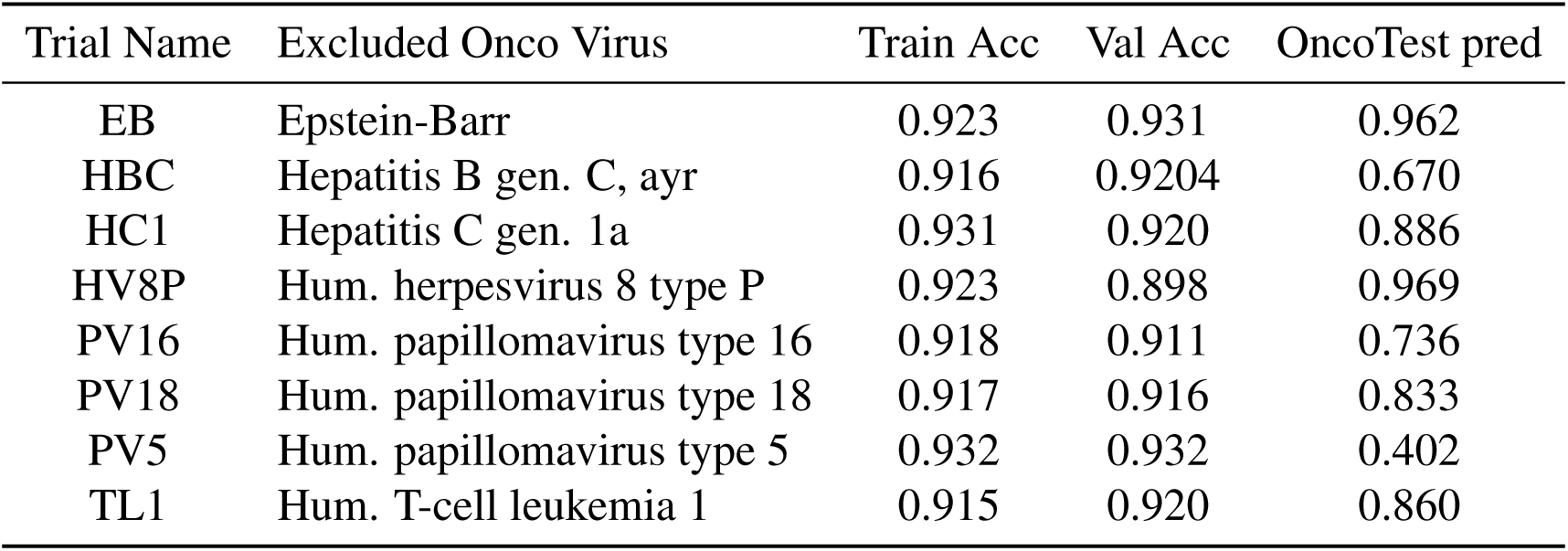
The table presents the performance results of individual Perceptron models trained using datasets in which samples containing specific oncogenic viruses were excluded, specifically the training and validation accuracy at the end of each model training. The *OncoTest pred* column contains the accuracy values of the predictions performed over the samples containing the excluded oncogenic virus PPI layer.

Following the training of the perceptron, it is possible to extract information about relevant input features, in this case, human proteins, which can be associated with classification into the oncogenic or non-oncogenic classes. This is achieved by analyzing the values of the perceptron’s weights. These weights are organized into two sets, each corresponding to a possible output, corresponding to whether the sample contains an oncogenic layer or not. Each set has a number of entries equal to the number of human proteins. The concept is that if the absolute difference between the values of two weights corresponding to the same protein but associated with the two possible outputs is significant, it can be inferred that when a protein is characterized by substantial centrality in the network, it is strongly positively associated with one output and strongly negatively associated with the other.

Employing this method and intersecting the top 200 proteins obtained from each training trial of the perceptron, a list of 81 proteins is generated. By performing a functional enrichment analysis based on Gene Ontology, the results show also in this case a great relevance of processes involved in chromatin structure. The complete list of such proteins and the table depicting the result of the Gene Ontology functional enrichment analysis are reported in the S9 Appendix.

### Specificity on oncogenic non-oncogenic classification task

Viruses can be classified upon a large number of features, and in general they share many characteristics between each other. Oncogenicity is only one of them, and arguably not even one of the most distinctive for viruses classification. Taking also into account the number of parameters the perceptron model has, it is reasonable to assume that it would be able to perform a classification similar to the one described before, even for a different set of target viruses (in the previous section this target viruses set corresponded to the oncogenic ones). To do so we considered a new class of target viruses, which were chosen randomly with the constraints of having virus-host interaction network sizes distribution is similar to the one of oncogenic viruses. In such a way such viruses do not constitute a class by themselves a priori. The dataset composed by multilayers, containing or not one layer corresponding to a “target virus”, is built, and the perceptron is trained with multiple trials as described before. The performance was very good also in this case, and the functional enrichment analysis performed over the most representative proteins for the classification (in an analogous way as described in the previous paragraph) reveals a large number of highly enriched pathways. But such pathways are not anymore related to chromatin structure as in the case of the classification oncogenic/non-oncogenic, revealing that the machine learning procedure can be generalized also to other classification tasks, but the results obtained for the case study can still be considered meaningful and taken as valuable insights on the subject. More details about this results, including the ones related to GO functional enrichment analysis, can be found in S6 Appendix, S10 Appendix.

## Discussion

Studying oncogenicity in viruses is complex, since viruses employ many different mechanisms to infect their hosts. This study aims to gain insights into these topics using a network medicine approach based on Protein-Protein Interaction (PPI) networks, with the goal of offering valuable information for ongoing medical research on viral oncogenicity.

The multilayer framework has shown significant potential for this analysis, enabling precise and quantitative descriptions of the impact that specific viruses can have on the human interactome. Topological analysis reveals clear statistical differences between samples obtained from oncogenic and non-oncogenic virus PPI networks. This distinction is further supported by the combined feature analysis, where the separation between the two categories becomes even more pronounced.

Focusing on the distribution of the Largest Intersected Component (LIC) and Largest Viable Component (LVC) dimensions, higher values associated with oncogenic samples suggest that oncogenic viruses generally target human proteomic regions that are more similar to each other compared to non-oncogenic viruses. This may be linked to specific mechanisms associated with oncogenicity. To explore this possibility, we extracted the set of nodes that are connected in all the PPI networks associated with oncogenic viruses. Subsequently, we performed an enrichment analysis with Gene Ontology, considering molecular functions, biological processes and cellular components. Interestingly, many results are related to proteins involved in chromatin structure, a fundamental aspect of cell life, which influences many of its processes[69]. In particular, alterations in chromatin structure have been found to contribute to the development and progression of cancer[70]. Various processes accompany alterations in chromatin organization in cancer cells, including changes in DNA methylation patterns, histone modifications, and higher-order chromatin folding [71][72].

One key aspect is the epigenetic regulation of gene expression through modifications of chromatin structure. Epigenetic modifications, including DNA methylation and histone modifications, can lead to the silencing or activation of specific genes involved in cancer-related processes such as cell proliferation, differentiation, and metastasis.

Considering the distribution of the percolation critical point, a statistically significant difference between oncogenic viruses and others is evident. Specifically, samples composed of oncogenic virus layers tend to have a higher associated critical point, indicating that the system is more robust against targeted attacks. This implies that this kind of viruses may target areas of the human interactome that are more robustly connected, potentially because they belong to more important and structured processes within human cells, likely of earlier evolutionary origin and relevant to a broader number of cell types.

As for quantities related to the community partition of the system, a marked difference exists between oncogenic and non-oncogenic cases. The oncogenic case is characterized by a larger number of modules and lower modularity than the non-oncogenic case. This suggests that it may be more challenging to identify clear community structures in oncogenic systems due to a more homogeneous distribution of links between nodes, an effect that could be related to a more systemic nature of oncongenic viruses.

While each of these measures offers insights into the separation of the two virus categories, no clear, easily identifiable threshold between them is observed. To address this, we propose integrating different features to improve the classification procedure. UMAP visualization effectively illustrates this, with a clear separation between oncogenic and non-oncogenic regions. As the number of oncogenic layers increases, the distribution shifts from the oncogenic region to the non-oncogenic one, enforcing the confidence in the classification results.

It is well known that the data from which the models used for the analysis are built are characterized by a huge degree of uncertainty and potentially huge biases due to experimental reasons[73, 74]. For this reason we tested our results by randomizing the model, which consisted in performing random rewiring of the complete human interactome from which each time the virus-host interaction networks were extracted. Similar analysis to the ones described above were performed to check for statistical differences that could support the assumption according to which the original data can indeed form a model able to well describe some aspect of the human cell biology. The complete results of such analysis are reported in S2 Appendix, S3 Appendix, S4 Appendix, S5 Appendix, and they all lead to the conclusion that the randomized models indeed do not provide highly relevant biological results, reinforcing the results reported in the article.

The next step involves considering a global description of each network which is used to propose a method to perform classification with the ultimate goal to isolate the contribution of the single layer, and at the same time also being able to keep track of the relevance of the single nodes. We introduced a machine learning pipeline based on the perceptron architecture, which allows for the extraction of a set of relevant nodes for classification by analyzing the weights, as detailed in the results section. By performing GO functional enrichment analysis over this set, we obtain results very similar to the ones obtained from the largest components analysis, consistent with our initial findings.

Upon completing perceptron training with different datasets, we apply a procedure to extract the most relevant input features for classification. The top 200 proteins from each trial are intersected to obtain a core set of 81 key proteins. We then use this core to perform a Gene Ontology molecular function enrichment analysis to explore potential associations between these proteins and cancer. The results reveal a high relevance of processes involved in chromatin structure, confirming the insights coming from the previous analysis.

A critical consideration is the potential presence of biases that could impact the final classification results. Viruses can be classified upon various characteristics, such as the type of nucleic acid they contain (DNA or RNA), the type of infection they cause in the body (intestinal, respiratory…), size and morphology (helical, icosahedral, prolate, enveloped,…) and taxonomic identification. The categorisation of DNA vs. RNA viruses was explored, in particular regarding the results of the machine learning method -as illustrated in S7 Appendix-, and appears not to be a relevant bias factor in the classification task. The other possible differentiation between viruses could be explored in future work, leading to a more robust classification analysis.

Another possible future improvement lies in refining the virus-host PPI network construction. Rather than simply subsetting the human PPI network to the first neighbors of virus-targeted proteins, future approaches could integrate a broader biological context, including indirect interaction pathways, structural proximity, or additional data sources to achieve a more accurate representation of viral influence on host cellular processes. Another potential enhancement involves quantifying oncogenicity as a spectrum rather than a binary classification, which could improve sensitivity to partially oncogenic viruses. Further, exploring virus characteristics—such as infection type or taxonomic data—could make the classification model more robust and biologically informative.

Overall, our findings suggest that it is possible to employ a procedure to inspect possible long term effects of viruses on humans without using data coming from clinical research. This can be applied to novel viruses, providing as example some insights that could help to orientate clinical research.

## Acknowledgments

F.Z. is supported by a Decreto Ministeriale n. 118 del 02/03/2023, M4C1 I. 4.1 from Piano Nazionale di Ripresa e Resilienza (PNRR) grant. V.P. is supported by INSERM, the Fondation Toulouse Cancer Santé and Pierre Fabre Research Institute as part of the Chair of Bioinformatics in Oncology of the CRCT.

## Author contributions

M.D.D., V.P. and F.Z. designed research; M.D.D., V.P. and F.Z. performed research; F.Z. analyzed data; and M.D.D., V.P. and F.Z. wrote the paper

## Competing interests

The authors declare no competing interests.

## Availability of data and materials

This study utilizes the dataset described in [48], where additional information and resources can be found. The jupyter notebooks used for the analysis and the figure production can be found in https://github.com/francescozambelli/OncoVirus.

## APPENDIX: Materials and methods

### Dataset overview

The study relies on BIOSTRING [28], a dataset created by integrating data from STRING and BIOGRID aimed to describe the human interactome and its interactions with proteins belonging to a 80 different viruses. For both human-human and virus-human PPIs, we chose to consider a threshold equal to 0.7 on the confidence score provided by STRING and BIOGRID. This results in a huge variety of interaction types, including genetic, co-expression, physical, gene neighborhood, gene fusion, gene homology, text mining links, giving our model a broad descriptive view of the underlying systems. Following this procedure the human interactome consists in a network of 15,131 nodes and 719,552 edges. The number of virus-host interactions depends on the single virus, and further details can be found in [28]. The classification of viruses into oncogenic and non-oncogenic categories was established on strongly confirmed clinical literature.

### Mathematical model for virus-host interaction networks

The construction of virus-human interaction PPI networks is graphically described in the top section of Figure 1. The starting point consists in the entire human PPI network *G_h_* = (*V_h_, E_h_*) in which nodes *V_h_* correspond to human proteins and the edges *E_h_* represent functional protein interactions from BIOSTRING. As described in the results sections, for virus-host interactions the focus here is on identifying the human interactome regions most likely to be affected by the interactions with viral proteins.

To implement this approach, we begin by considering the subset of proteins in the entire human PPI network that are documented to interact with viral proteins. Restricting the network to only these nodes often results in a network that is too small and dispersed, offering limited information. To address this, we expand our focus to include another set of proteins that are likely to be influenced by the interaction with viral proteins, specifically the nearest neighbors of the directly targeted proteins.

At this stage, for each virus *v_i_*, *i* = 1*, …, N_v_*, with *N_v_* equal to the number of viruses for which PPI data are present in the dataset, we can construct a dedicated PPI network *G_i_* = (*V_i_, E_i_*) in which the set of nodes *V_i_* consists in the merged set of proteins directly targeted and their closest neighbors from the broader human PPI network, while the edge set *E_i_* contains all the human PPI interaction links involving at least one of the proteins in the *V_i_* set.

### Multilayer networks formalism and multi-pagerank versatility

The mathematical formulation of multilayer networks involves representing these systems as a collection of nodes and edges across multiple layers. Each layer describe how the same set of nodes interact in different conditions of the system. Specifically, such multilayer networks consist in a set of *N* physical nodes, repeated in each of the *L* layers composing the systems, and connected in each layer through a different set of edges *E_l_*, *l* = 1*, …, L*.

As regards the mathematical formalism, multilayer networks appears to fit particularly well with tensorial formulation[59]. In particular it is possible to define a multilinear object in the space ℝ*^NxLxLxN^*, which corresponds to the multi-adjacency tensor describing both inter- and intra-layer edges:

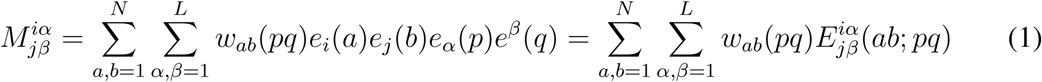

where *e_i_*(*a*) and *e^j^*(*b*) are the covariant and contravariant canonical rank-1 tensors, and *w_ab_*(*pq*) encodes the strength of the interactions between node *a* in layer *p* and node *b* in layer *q*.

Through the work, we used two different types of multilayer structure to represent the data, depending on the specific analysis to be performed. The first one corresponds to all-interconnected multilayer structure, in which each replica of each node is connected directly with all the others. Such structure is used as example when considering the components analysis and the percolation. The second one is the edge-colored framework, in which the inter-layer links are not specified. Such model is used for the multi-pagerank centrality computation, which would strongly depend on the inter-layer links strengths if those were specified.

The multi-pagerank versatility measure is defined as the asymptotic occupation probability of a given node by a random walker that can travel through nodes of the different layers subjected to teleportation [59, 60]. Mathematically, we start by defining a transition tensor 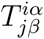, which gives the probability to transit from a node *i* in layer *α* to a node *j* in layer *β*. Such probabilities are proportional to the edge strengths between the two nodes, and the sum of the probabilities to go from a specific node to all the others should sum up to 1, as per the definition of probability. Once the transition tensor is defined, by solving the eigenvalue problem it is possible to find the set of stationary occupancy probabilities which correspond to the value entries of the eigenvector corresponding to eigenvalue 1.

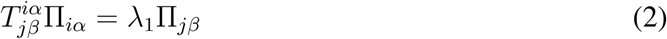

The probabilities Π*_iα_* define the random walk occupation centralities, which can be further compressed by aggregating values corresponding to replicas of the same node, thus obtaining a vector of scores for each physical node. The multi-pagerank versatility derives from a variant of such measure. In fact the transition tensor can be modified to incorporate a term corresponding to a uniform probability to transit to each node in each one of the layers of the network.

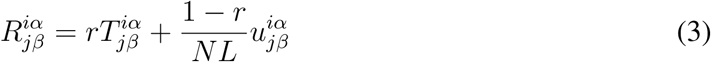

with *N* the number of physical nodes, *L* the number of layers, 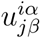 the rank-4 tensor with all the entries equal to 1, and *r* the coefficient which determines the probability of teleportation at each step. By solving the same eigenvalue problem and appropriately aggregating the eigenvector entries, it is possible to obtain the occupation probabilities associated to this new random walk process. In this work the *r* coefficient is set to 0.85, following examples from literature and computational implementations[60, 75].

### Random rewiring with configuration model

The rewiring of the human interactome used for proposing a null model over which to test the results from the analysis is performed by using the configuration model. It randomly rewires the graph by maintaining the degree sequence of the nodes unaltered. In this way the number of edges for each node is preserved, which is a desirable feature if we want to create a topology not too dissimilar, and in this way reasonably comparable, with the original one.

### Functional enrichment analysis

To quantify to what extent a list of proteins might indicate the involvement of a biological pathway, function or process, we performed functional enrichment analysis comprising Gene Ontology molecular function, biological processes, and cellular components. The tool used is ToppGene(https://toppgene.cchmc.org/), and the p-value threshold over which entries were considered to be significant is chosen as 5 · 10^−2^ considering the FDR correction.

## Supporting information

**S1 Appendix Components analysis, LCC.**

**S2 Appendix. Comparison with null model.**

**S3 Appendix. Comparison with null model, LVC.**

**S4 Appendix. Comparison with null model, percolation.**

**S5 Appendix. Comparison with null model, community partition.**

**S6 Appendix. Machine Learning cross-checking.**

**S7 Appendix. DNA vs RNA viruses.**

**S8 Appendix. GO enrichment analysis results, common nodes between all oncogenic layers.**

**S9 Appendix. GO enrichment analysis results, highly relevant proteins from perceptron weights analysis.**

**S10 Appendix. GO enrichment analysis results, highly relevant proteins from perceptron weights analysis with different target set.**

## Notes

https://github.com/francescozambelli/OncoVirus

### Competing Interest Statement

The authors have declared no competing interest.

## References and Notes

[1] Hughes, J. M., Wilson, M. E. & Sejvar, J. J. The Long-Term Outcomes of Human West Nile Virus Infection. Clinical Infectious Diseases 44, 1617–1624 (2007). URL 10.1086/518281.

[2] Kneyber, M., Steyerberg, E., de Groot, R. & Moll, H. Long-term effects of respiratory syncytial virus (RSV) bronchiolitis in infants and young children: a quantitative review. Acta Paediatrica 89, 654–660 (2000). URL https://onlinelibrary.wiley.com/doi/abs/10.1111/j.1651-2227.2000.tb00359.x. eprint: https://onlinelibrary.wiley.com/doi/pdf/10.1111/j.1651-2227.2000.tb00359.x.

[3] Clutton, G., et al. The differential short- and long-term effects of HIV-1 latency-reversing agents on T cell function. Scientific Reports 6, 30749 (2016). URL https://www.nature.com/articles/srep30749. Number: 1 Publisher: Nature Publishing Group.

[4] Dunmire, S. K., Verghese, P. S. & Balfour, H. H. Primary Epstein-Barr virus infection. Journal of Clinical Virology 102, 84–92 (2018). URL https://www.sciencedirect.com/science/article/pii/S1386653218300635.

[5] Nguyen, H.-N. T. et al. SARS-CoV-2 m protein facilitates malignant transformation of breast cancer cells. Frontiers in Oncology 12 (2022). URL 10.3389/fonc.2022.923467.

[6] Gómez-Carballa, A., Martinón-Torres, F. & Salas, A. Is SARS-CoV-2 an oncogenic virus? Journal of Infection 85, 573–607 (2022). URL 10.1016/j.jinf.2022.08.005.

[7] Costanzo, M., Giglio, M. A. R. D. & Roviello, G. N. Deciphering the relationship between SARS-CoV-2 and cancer. International Journal of Molecular Sciences 24, 7803 (2023). URL 10.3390/ijms24097803.

[8] Schiller, J. T. & Lowy, D. R. Virus infection and human cancer: An overview. In Viruses and Human Cancer, 1–10 (Springer Berlin Heidelberg, 2013). URL 10.1007/978-3-642-38965-8_1.

[9] Thompson, M. P. & Kurzrock, R. Epstein-barr virus and cancer. Clinical Cancer Research 10, 803–821 (2004). URL 10.1158/1078-0432.ccr-0670-3.

[10] Gioia, U. et al. Sars-cov-2 infection induces dna damage, through chk1 degradation and impaired 53bp1 recruitment, and cellular senescence. Nature Cell Biology 25, 550–564 (2023). URL 10.1038/s41556-023-01096-x.

[11] Barabási, A.-L., Gulbahce, N. & Loscalzo, J. Network medicine: a network-based approach to human disease. Nature Reviews Genetics 12, 56–68 (2010). URL 10.1038/nrg2918.

[12] Ivanov, P. C., Liu, K. K. L. & Bartsch, R. P. Focus on the emerging new fields of network physiology and network medicine. New Journal of Physics 18, 100201 (2016). URL 10.1088/1367-2630/18/10/100201.

[13] Zhou, X., Menche, J., Barabási, A.-L. & Sharma, A. Human symptoms–disease network. Nature Communications 5 (2014). URL 10.1038/ncomms5212.

[14] Lee, L. Y.-H. & Loscalzo, J. Network medicine in pathobiology. The American Journal of Pathology 189, 1311–1326 (2019). URL 10.1016/j.ajpath.2019.03.009.

[15] Goh, K.-I. et al. The human disease network. Proceedings of the National Academy of Sciences 104, 8685–8690 (2007).

[16] Halu, A., Domenico, M. D., Arenas, A. & Sharma, A. The multiplex network of human diseases. npj Systems Biology and Applications 5 (2019). URL 10.1038/s41540-019-0092-5.

[17] Li, S. et al. Comprehensive characterization of human–virus protein-protein interactions reveals disease comorbidities and potential antiviral drugs. Computational and Structural Biotechnology Journal 20, 1244–1253 (2022). URL 10.1016/j.csbj.2022.03.002.

[18] Gysi, D. M. et al. Network medicine framework for identifying drug-repurposing opportunities for COVID-19. Proceedings of the National Academy of Sciences 118 (2021). URL 10.1073/pnas.2025581118.

[19] Rual, J.-F. et al. Towards a proteome-scale map of the human protein–protein interaction network. Nature 437, 1173–1178 (2005). URL 10.1038/nature04209.

[20] De Las Rivas, J. & Fontanillo, C. Protein-protein interaction networks: unraveling the wiring of molecular machines within the cell. Briefings in Functional Genomics 11, 489–496 (2012). URL 10.1093/bfgp/els036.

[21] Braun, P. & Gingras, A. History of protein–protein interactions: From egg-white to complex networks. PROTEOMICS 12, 1478–1498 (2012). URL 10.1002/pmic.201100563.

[22] Szklarczyk, D. et al. STRING v11: protein–protein association networks with increased coverage, supporting functional discovery in genome-wide experimental datasets. Nucleic Acids Research 47, D607–D613 (2018). URL 10.1093/nar/gky1131.

[23] Choobdar, S. et al. Assessment of network module identification across complex diseases. Nature methods 16, 843–852 (2019).

[24] Srinivasan, S. et al. Structural genomics of sars-cov-2 indicates evolutionary conserved functional regions of viral proteins. Viruses 12, 360 (2020). URL 10.3390/v12040360.

[25] Gordon, D. E. et al. A SARS-CoV-2 protein interaction map reveals targets for drug repurposing. Nature 583, 459–468 (2020). URL 10.1038/s41586-020-2286-9.

[26] Zhou, W., Chen, Z., Fang, Z. & Xu, D. Network analysis for elucidating the mechanisms of shenfu injection in preventing and treating covid-19 combined with heart failure. Computers in Biology and Medicine 148, 105845 (2022). URL 10.1016/j.compbiomed.2022.105845.

[27] Alzahrani, F. A., Khan, M. F. & Ahmad, V. Recognition of differentially expressed molecular signatures and pathways associated with covid-19 poor prognosis in glioblastoma patients. International Journal of Molecular Sciences 24, 3562 (2023). URL 10.3390/ijms24043562.

[28] Verstraete, N. et al. CovMulNet19, integrating proteins, diseases, drugs, and symptoms: A network medicine approach to COVID-19. Network and Systems Medicine 3, 130–141 (2020). URL 10.1089/nsm.2020.0011.

[29] Ding, Q. & Zhao, H. Long-term effects of SARS-CoV-2 infection on human brain and memory. Cell Death Discovery 9 (2023). URL 10.1038/s41420-023-01512-z.

[30] Al-Aly, Z. & Topol, E. Solving the puzzle of long covid. Science 383, 830–832 (2024). URL 10.1126/science.adl0867.

[31] Klein, J. et al. Distinguishing features of long covid identified through immune profiling. Nature 623, 139–148 (2023). URL 10.1038/s41586-023-06651-y.

[32] De Domenico, M. et al. Mathematical formulation of multilayer networks. Physical Review X 3, 041022 (2013).

[33] Domenico, M. D. More is different in real-world multilayer networks. Nature Physics 19, 1247–1262 (2023). URL 10.1038/s41567-023-02132-1.

[34] Nicosia, V., Bianconi, G., Latora, V. & Barthelemy, M. Growing multiplex networks. Physical review letters 111, 058701 (2013).

[35] Battiston, F., Nicosia, V. & Latora, V. The new challenges of multiplex networks: Measures and models. The European Physical Journal Special Topics 226, 401–416 (2017).

[36] Cardillo, A. et al. Emergence of network features from multiplexity. Scientific reports 3, 1344 (2013).

[37] Gallotti, R., Sacco, P. & Domenico, M. D. Complex urban systems: Challenges and integrated solutions for the sustainability and resilience of cities. Complexity 2021, 1–15 (2021). URL 10.1155/2021/1782354.

[38] Pilosof, S., Porter, M. A., Pascual, M. & Kéfi, S. The multilayer nature of ecological networks. Nature Ecology and Evolution 1 (2017). URL 10.1038/s41559-017-0101.

[39] Wang, Z., Wang, L., Szolnoki, A. & Perc, M. Evolutionary games on multilayer networks: a colloquium. The European physical journal B 88, 1–15 (2015).

[40] Bargigli, L., di Iasio, G., Infante, L., Lillo, F. & Pierobon, F. The multiplex structure of interbank networks. Quantitative Finance 15 (2014). URL 10.1080/14697688.2014.968356.

[41] Buldyrev, S. V., Parshani, R., Paul, G., Stanley, H. E. & Havlin, S. Catastrophic cascade of failures in interdependent networks. Nature 464, 1025–1028 (2010).

[42] Granell, C., Gómez, S. & Arenas, A. Dynamical interplay between awareness and epidemic spreading in multiplex networks. Physical Review Letters 111 (2013). URL 10.1103/physrevlett.111.128701.

[43] Radicchi, F. & Bianconi, G. Redundant interdependencies boost the robustness of multiplex networks. Physical Review X 7, 011013 (2017).

[44] Jeude, J. v. L. d., Aste, T. & Caldarelli, G. The multilayer structure of corporate networks. New Journal of Physics 21, 025002 (2019).

[45] Wang, M., Sunkel, B. D., Ray, W. C. & Stanton, B. Z. Chromatin structure in cancer. BMC Molecular and Cell Biology 23 (2022). URL 10.1186/s12860-022-00433-6.

[46] Zhao, S., Allis, C. D. & Wang, G. G. The language of chromatin modification in human cancers. Nature Reviews Cancer 21, 413–430 (2021). URL 10.1038/s41568-021-00357-x.

[47] Akdemir, K. C. et al. Disruption of chromatin folding domains by somatic genomic rearrangements in human cancer. Nature Genetics 52, 294–305 (2020). URL 10.1038/s41588-019-0564-y.

[48] Ghavasieh, A., Bontorin, S., Artime, O., Verstraete, N. & Domenico, M. D. Multiscale statistical physics of the pan-viral interactome unravels the systemic nature of SARS-CoV-2 infections. Communications Physics 4 (2021). URL 10.1038/s42005-021-00582-8.

[49] Oughtred, R., Rust, J., Chang, C. & al. The biogrid database: A comprehensive biomedical resource of curated protein, genetic, and chemical interactions. Protein Science 30, 187–200 (2021). URL 10.1002/pro.3978.

[50] Szklarczyk, D., et al. STRING v10: protein–protein interaction networks, integrated over the tree of life. Nucleic Acids Research 43, D447–D452 (2014). URL 10.1093/nar/gku1003.

[51] Cook, H., Doncheva, N., Szklarczyk, D., von Mering, C. & Jensen, L. Viruses.string: A virus-host protein-protein interaction database. Viruses 10 (2018).

[52] Brito, A. F. & Pinney, J. W. Protein–protein interactions in virus–host systems. Frontiers in Microbiology 8 (2017). URL 10.3389/fmicb.2017.01557.

[53] Yang, S., Fu, C., Lian, X., Dong, X. & Zhang, Z. Understanding human-virus protein-protein interactions using a human protein complex-based analysis framework. mSystems 4 (2019). URL 10.1128/msystems.00303-18.

[54] Baxter, G. J., Dorogovtsev, S. N., Goltsev, A. V. & Mendes, J. F. F. Avalanche collapse of interdependent networks. Physical Review Letters 109 (2012). URL 10.1103/physrevlett.109.248701.

[55] Azimi-Tafreshi, N., Gómez-Gardeñes, J. & Dorogovtsev, S. N. k-core percolation on multiplex networks. Physical Review E 90 (2014). URL 10.1103/physreve.90.032816.

[56] Grassberger, P. Percolation transitions in the survival of interdependent agents on multiplex networks, catastrophic cascades, and solid-on-solid surface growth. Physical Review E 91 (2015). URL 10.1103/physreve.91.062806.

[57] Stella, M., Beckage, N. M., Brede, M. & Domenico, M. D. Multiplex model of mental lexicon reveals explosive learning in humans. Scientific Reports 8 (2018). URL 10.1038/s41598-018-20730-5.

[58] Artime, O. et al. Robustness and resilience of complex networks. Nature Reviews Physics 1–18 (2024).

[59] De Domenico, M. et al. Mathematical formulation of multilayer networks. Phys. Rev. X 3, 041022 (2013). URL https://link.aps.org/doi/10.1103/PhysRevX.3.041022.

[60] De Domenico, M., Solé-Ribalta, A., Omodei, E., Gómez, S. & Arenas, A. Ranking in interconnected multilayer networks reveals versatile nodes. Nature Communications 6 (2015). URL 10.1038/ncomms7868.

[61] Fortunato, S. Community detection in graphs. Physics Reports 486, 75–174 (2010). URL 10.1016/j.physrep.2009.11.002.

[62] Newman, M. E. J. Communities, modules and large-scale structure in networks. Nature Physics 8, 25–31 (2011). URL 10.1038/nphys2162.

[63] Newman, M. E. Modularity and community structure in networks. Proceedings of the national academy of sciences 103, 8577–8582 (2006).

[64] Blondel, V. D., Guillaume, J.-L., Lambiotte, R. & Lefebvre, E. Fast unfolding of communities in large networks. Journal of Statistical Mechanics: Theory and Experiment 2008, P10008 (2008). URL 10.1088/1742-5468/2008/10/ P10008.

[65] Peixoto, T. P. Bayesian stochastic blockmodeling (2019). URL 10.1002%2F9781119483298.ch11.

[66] Peixoto, T. P. Inferring the mesoscale structure of layered, edge-valued, and time-varying networks. Physical Review E 92 (2015). URL 10.1103/physreve.92.042807.

[67] Spirin, V. & Mirny, L. A. Protein complexes and functional modules in molecular networks. Proceedings of the National Academy of Sciences 100, 12123–12128 (2003). URL 10.1073/pnas.2032324100.

[68] Newman, M. E. J. Modularity and community structure in networks. Proceedings of the National Academy of Sciences 103, 8577–8582 (2006). URL 10.1073/pnas.0601602103.

[69] Pancaldi, V. Network models of chromatin structure. Current Opinion in Genetics & Development 80, 102051 (2023). URL 10.1016/j.gde.2023.102051.

[70] Wang, M., Sunkel, B. D., Ray, W. C. & Stanton, B. Z. Chromatin structure in cancer. BMC Molecular and Cell Biology 23 (2022). URL 10.1186/s12860-022-00433-6.

[71] Zhao, S., Allis, C. D. & Wang, G. G. The language of chromatin modification in human cancers. Nature Reviews Cancer 21, 413–430 (2021). URL 10.1038/s41568-021-00357-x.

[72] Brock, M. V., Herman, J. G. & Baylin, S. B. Cancer as a manifestation of aberrant chromatin structure. The Cancer Journal 13, 3–8 (2007). URL 10.1097/ppo.0b013e31803c5415.

[73] Wodak, S. J., Pu, S., Vlasblom, J. & Seéraphin, B. Challenges and rewards of interaction proteomics. Molecular & Cellular Proteomics 8, 3–18 (2009). URL 10.1074/mcp.R800014-MCP200.

[74] Peel, L., Peixoto, T. P. & De Domenico, M. Statistical inference links data and theory in network science. Nature Communications 13 (2022). URL 10.1038/s41467-022-34267-9.

[75] De Domenico, M., Porter, M. A. & Arenas, A. Muxviz: a tool for multilayer analysis and visualization of networks. Journal of Complex Networks 3, 159–176 (2014). URL 10.1093/comnet/cnu038.

