## Supplemental Info for "Unraveling the network signatures of oncogenicity in virus-human protein-protein interactions"

```
File: main.tex
Encoding: utf8
Sum count: 4074
Words in text: 3132
Words in headers: 90
Words outside text (captions, etc.): 810
Number of headers: 14
Number of floats/tables/figures: 15
Number of math inlines: 39
Number of math displayed: 3
Subcounts:
  text+headers+captions (#headers/#floats/#inlines/#displayed)
  1+10+0 (1/0/0/0) _top_
  233+6+0 (1/0/10/0) Section: Mathematical model for virus-host interactio
  505+6+0 (1/0/24/3) Section: Multilayer networks formalism and multi-page
  162+3+0 (1/0/4/0) Section: Components analysis, LCC
  222+4+0 (1/0/0/0) Section: Comparison with null model
  424+5+0 (1/0/0/0) Section: Comparison with null model, LVC
  153+2+0 (1/0/0/0) Subsection: Enrichment analysis
  181+5+0 (1/0/0/0) Section: Comparison with null model, percolation
  56+6+0 (1/0/0/0) Section: Comparison with null model, community partitio
  287+3+0 (1/0/0/0) Section: Machine Learning cross-checking
  443+4+0 (1/0/1/0) Section: DNA vs RNA viruses
  85+10+0 (1/0/0/0) Section: GO enrichment analysis results, common nodes b
  142+11+0 (1/0/0/0) Section: GO enrichment analysis results, highly releva
  238+15+810 (1/15/0/0) Section: GO enrichment analysis results, highly rel

File: output.bbl
Encoding: utf8
Sum count: 0
Words in text: 0
Words in headers: 0
Words outside text (captions, etc.): 0
Number of headers: 0
Number of floats/tables/figures: 0
Number of math inlines: 0
Number of math displayed: 0
```

### Unraveling the network signatures of oncogenicity in virus-human protein-protein interactions

Francesco Zambelli[1], Vera Pancaldi [2,3], Manlio De Domenico [1,4,5,*]

University of Padova, 35123 Padova, Italy University of Padova, Padova, Italy
[2]Centre de Recherches en Cancérologie de Toulouse (CRCT), UMR1037 Inserm, ERL5294 CNRS, Toulouse, France
[3]University Paul Sabatier III, Toulouse, France
[4]Padua Center for Network Medicine, University of Padua, University of Padova, Padova, Italy
[5]Istituto Nazionale di Fisica Nucleare, Sez. Padova, Italy

#### 1 Mathematical model for virus-host interaction networks

The construction of virus-human interaction PPI networks is graphically described in the top section of Figure 1 of the main article. The starting point consists in the entire human PPI network $G_h = (V_h, E_h)$ in which nodes $V_h$ correspond to human proteins and the edges $E_h$ represent functional protein interactions from BIOSTRING. As described in the results sections, for virus-host interactions the focus here is on identifying the human interactome regions most likely to be affected by the interactions with viral proteins.

To implement this approach, we begin by considering the subset of proteins in the entire human PPI network that are documented to interact with viral proteins. Restricting the network to only these nodes often results in a network that is too small and dispersed, offering limited information. To address this, we expand our focus to include another set of proteins that are likely to be influenced by the interaction with viral proteins, specifically the nearest neighbors of the directly targeted proteins.

At this stage, for each virus $v_i$, $i = 1, ..., N_v$, with $N_v$ equal to the number of viruses for which PPI data are present in the dataset, we can construct a dedicated PPI network $G_i = (V_i, E_i)$ in which the set of nodes $V_i$ consists in the merged set of proteins directly targeted and their closest neighbors from the broader human PPI network, while the edge set $E_i$ contains all the human PPI interaction links involving at least one of the proteins in the $V_i$ set.

### 2 Multilayer networks formalism and multi-pagerank versatility

The mathematical formulation of multilayer networks involves representing these systems as a collection of nodes and edges across multiple layers. Each layer describe how the same set of nodes interact in different conditions of the system. Specifically, such multilayer networks consist in a set of $N$ physical nodes, repeated in each of the $L$ layers composing the systems, and connected in each layer through a different set of edges $E_l$, $l = 1, ..., L$.

As regards the mathematical formalism, multilayer networks appears to fit particularly well with tensorial formulation[1]. In particular it is possible to define a multilinear object in the space $\mathbb{R}^{NxLxLxN}$, which corresponds to the multi-adjacency tensor describing both inter- and intra-layer edges:

$$M^{i\alpha}_{j\beta} = \sum_{a,b=1}^{N} \sum_{\alpha,\beta=1}^{L} w_{ab}(pq) e_i(a) e_j(b) e_\alpha(p) e^\beta(q) = \sum_{a,b=1}^{N} \sum_{\alpha,\beta=1}^{L} w_{ab}(pq) E^{i\alpha}_{j\beta}(ab; pq) \quad (1)$$

where $e_i(a)$ and $e^j(b)$ are the covariant and contravariant canonical rank-1 tensors, and $w_{ab}(pq)$ encodes the strength of the interactions between node $a$ in layer $p$ and node $b$ in layer $q$.

Through the work, we used two different types of multilayer structure to represent the data, depending on the specific analysis to be performed. The first one corresponds to all-interconnected multilayer structure, in which each replica of each node is connected directly with all the others. Such structure is used as example when considering the components analysis and the percolation. The second one is the edge-colored framework, in which the inter-layer links are not specified. Such model is used for the multi-pagerank centrality computation, which would strongly depend on the inter-layer links strengths if those were specified.

The multi-pagerank versatility measure is defined as the asymptotic occupation probability of a given node by a random walker that can travel through nodes of the different layers subjected to teleportation [1, 2]. Mathematically, we start by defining a transition tensor $T^{i\alpha}_{j\beta}$, which gives the probability to transit from a node $i$ in layer $\alpha$ to a node $j$ in layer $\beta$. Such probabilities are proportional to the edge strengths between the two nodes, and the sum of the probabilities to go from a specific node to all the others should sum up to 1, as per the definition of probability. Once the transition tensor is defined, by solving the eigenvalue problem it is possible to find the set of stationary occupancy probabilities which correspond to the value entries of the eigenvector corresponding to eigenvalue 1.

$$T^{i\alpha}_{j\beta}\Pi_{i\alpha} = \lambda_1 \Pi_{j\beta} \tag{2}$$

The probabilities $\Pi_{i\alpha}$ define the random walk occupation centralities, which can be further compressed by aggregating values corresponding to replicas of the same node, thus obtaining a vector of scores for each physical node. The multi-pagerank versatility derives from a variant of such measure. In fact the transition tensor can be modified to incorporate a term corresponding to a uniform probability to transit to each node in each one of the layers of the network.

$$R^{i\alpha}_{j\beta} = rT^{i\alpha}_{j\beta} + \frac{1-r}{NL}u^{i\alpha}_{j\beta} \tag{3}$$

with $N$ the number of physical nodes, $L$ the number of layers, $u^{i\alpha}_{j\beta}$ the rank-4 tensor with all the entries equal to 1, and $r$ the coefficient which determines the probability of teleportation at each step. By solving the same eigenvalue problem and appropriately aggregating the eigenvector entries, it is possible to obtain the occupation probabilities associated to this new random walk process. In this work the $r$ coefficient is set to $0.85$, following examples from literature and computational implementations[2, 3].

### 3 Components analysis, LCC

The analysis of the Largest Connected Components (LCCs) confirms that oncogenic PPI networks tend to be larger than non-oncogenic networks. This trend is evident when comparing the distributions of oncogenic ($O$) and non-oncogenic ($N$) networks in Fig.1(left). As more oncogenic layers are added to the multilayer networks, the distributions shift from $N$ to $O$.

Furthermore, it is evident that all the distributions are highly distinct from each other, as reinforced by observing the heatmap in Fig.1(right).

In conclusion, it can be observed that oncogenic viruses generally generate larger PPI networks compared to non-oncogenic viruses.

There are two possible explanations for this observation. First, it is possible that oncogenic viruses have a greater level of interaction with the human PPI network, resulting in larger virus-host PPI networks. Alternatively, this size disparity could be attributed to some form of bias. To address this hypothesis, the subsequent parts of the study will consider this factor and not solely rely on network size for the oncogenic/non-oncogenic classification.

### 4 Comparison with null model

In order to propose biologically plausible results, it is necessary to test the dataset against some kind of null model, to verify that it indeed contains biological information and the amount of unavoidable errors due to noisy experiments and data incompleteness, does not lead to a model with no biological correspondence.

This was done by using a randomization process, in which the underlying complete human PPI network gathered from online resources, from which all the virus-host interaction PPI networks are produced as described above, is randomly rewired. It is desirable to preserve the node degree sequence, in order to not fall to much far from the original model preserving some coarse graining properties but changing the actual specific edges, each one of those is supposed to have a specific biological counterpart in the cell. For this reason we chose a configurational model for the randomization.

After randomizing the complete human PPI network, it is possible to extract the virus-host interaction networks with the same procedure described before, by targeting the same proteins as described in the virus interaction dataset. By iterating this process it's possible to build multiple datasets by starting from different randomization of the complete network. At this point we compare some analysis results, which procedure is described in the article, also for this data, and we compare the results

### 5 Comparison with null model, LVC

The first analyzed quantity is the largest viable component size (LVC). For each of the virus-host interaction PPINs extracted form different randomization of the human PPIN, the size of the LVCs are computed for samples belonging from the combination sets as described in the *Identifying the core of nodes biologically relevant for oncogenesis* article section, and the comparison of the results between the null model and the original data are reported in fig.2.

It's possible to see that there is no evident discrepancy in the distribution of this quantity between the null model and the original data for neither of the combination sets. This can be explained by a simple reasoning about the number and kind of edges that are selected from the ones belonging to the complete human PPIN in the sub-setting process giving birth to virus-host interaction PPINs. This edges in fact can be of two types: either they can include a directly targeted node or they can connect two nearest neighbors of the same directly target node, or can be between two nearest neighbors of two different directly targeted nodes. The first number of the first kind of edges is preserved, given the configurational model and the fact that we consider the same directly targeted nodes. It is reasonable to think that the latter two edge types will be a second order effect, because they require a presence of a link between two regions of the network taken independently. This assumption could be verified by looking at the number of total edges differences between synthetic and true network for each virus (fig.4), and the number of edges which either the source of target node are directly targeted proteins(fig.3). In 3 it is of particular interest to see that the number of edges containing a directly targeted node are systematically higher in the synthetic networks than in the true network. More valuable information could be obtained by considering the distribution of such quantity with respect to the number of directly targeted proteins by a virus, as depicted in fig.6. It is firstly evident that in some cases the fraction of edges not containing a directly targeted protein are around 50-70%, but the totality of such cases happened with really small networks, where the number of directly targeted nodes is just some units. Another valuable insight comes from fig.5, where it's possible to observe a linear relation between the difference in number of edges containing a dir.targ. node between the synthetic and true networks, and the number of dir.targ. proteins by each virus.

#### 5.1 Enrichment analysis

Another possibility that could help to check if the randomization of the network indeed leads to a disruption of the biological content of the system, is looking at the functional enrichment analysis of the LVCs of the multilayers composed by the all 8 oncogenic viruses. It was chosen to consider Biological Processes, Molecular Functions and Cellular Components from Gene Ontology. The results are reported in Fig.7, in which each column is associated ot one of the GO categories, the rows correspond to values of FDR corrected p-value threshold under which we consider the item to be enriched. In each sub-figure, the blue histograms represent the distributions of the number of items enriched by the LVCs exctracted for 500 realization of the randomized model, while the red line represent the corresponding value for the original model. As expected, in all the cases the distributions associated to the randomized networks are strongly shifted towards 0.

### 6 Comparison with null model, percolation

The analysis of the LVC size distribution doesn't lead to a significant difference between the true and synthetic case, suggesting that relying on topological quantities relying on the static structure of the networks in not much informative under transformations performed by using the configurational model.

For this reason we take into consideration also the distribution of the critical point under the node percolation following the multi-pagerank ordering. This takes into account also how the information flows within the network, and is reasonable to think that a configurational model rewiring could lead to significant changes. This can be observed from fig.8, where is especially evident that the peak of the distributions for the different combination sets is significantly shifted between synthetic and true case. This straightens the assumption for which the network provided by the data indeed is not simply a random model. In particular the shift is towards higher values of the critical point for the synthetic network, indicating that such networks are less robust and end up in the disordered phase after the deletion of a smaller fraction of nodes.

### 7 Comparison with null model, community partition

As can be seen in fig.910, the results of the DCSBM community partition returns different results in the synthetic/randomized case with respect to the original network. It is particular meaningful the systematically smaller values for the modularity, indication that the partition into communities is less significant in the synthetic case compared to the original one.

### 8 Machine Learning cross-checking

A question could arise about the capabilities of the machine learning algorithm to perform the classification between networks associated to oncogenic and not-oncogenic viruses: what happens if we consider a different list of viruses as target set, i.e. the smaller group that we want to discriminate with respect to the others?

To do so we considered 8 not-oncogenic viruses, with network sizes similar to the ones of the oncogenic viruses set. In particular they are Human herpesvirus 1 strain 17, Human cytomegalovirus strain Merlin, Human herpesvirus 6A strain Uganda-1102, Human immunodeficiency virus type 1 group M, Yaba monkey tumor virus strain VR587, Varicella zoster virus strain Dumas, Saliviru A isolate Human-Nigeria NG J1-2007, Influenza B virus strain B-Lee-1940. We repeat the same analysis described in section *Machine learning classification* of the main article, this time considering the aforementioned 8 not-oncogenic viruses as belonging to the ”oncogenic class”, while the actual 8 oncogenic viruses are put in the ”not-oncogenic class”. There were performed multiple trials, each time taking out samples containing a layer associated to a virus belonging to the ”oncogenic class”. We considered the top 200 proteins whose absolute value of the difference between the weights of the two possible outputs were larger for each trial, and intersected them to obtain the final list. In tab.4 are presented the result of the Gene Ontology functional enrichment analysis. They show pathways highly enriched, which could be misleading at a first glance but becomes reasonable when thinking that the algorithm has as objective to spot up the features that could help to perform the task classification, which is not in contrast with the fact that the result presented in the article could actually describe common behaviours of oncogenic viruses.

### 9 DNA vs RNA viruses

In addition to the functional role of these specific proteins in tumor development, the difference in the nature of viruses may also contribute to the observed association. Viruses can be categorized as DNA viruses or RNA viruses. DNA viruses replicate their genetic material using host cellular machinery and DNA polymerases, primarily in the nucleus of the host cell. In contrast, RNA viruses replicate their genetic material using an RNA-dependent RNA polymerase (RdRp) and typically replicate in the cytoplasm of the host cell without accessing the nucleus.

Therefore, it is more likely that DNA viruses interact with histones, which are predominantly located in the nucleus of the cell. On the other hand, RNA viruses generally replicate in the cytoplasm, where histones are less abundant. However, it's important to note that certain RNA viruses, such as retroviruses, which convert their RNA genome into DNA intermediates, can interact with histones during the integration of viral DNA into the host cell's genome.

To further investigate this relationship, it is worth examining the distribution of viruses in the dataset between DNA and RNA viruses, including both oncogenic and non-oncogenic types. This analysis can reveal any imbalances or patterns. A schematic representation of the results is presented in Fig.11.

The number of DNA viruses is higher than RNA viruses in the oncogenic case, while in the non-oncogenic case, the trend is reversed. This discrepancy raises doubts about the classification task. The classifier may focus more on distinguishing between DNA and RNA viruses rather than between oncogenic and non-oncogenic viruses due to the imbalance in these classes.

Examining the performance of the perceptron and random forest classifiers (Tab.1, it is observed that the trial associated with the removal and testing of the *Human papillomavirus type 5* (a DNA virus) leads to the worst generalization performance.

This observation contradicts the hypothesis that the classification is based solely on virus type, as the performance should have significantly decreased when excluding and testing on an RNA virus. Furthermore, the training samples are even more imbalanced toward the DNA virus type, and the testing is performed with a virus of the opposite type.

Moreover, the performance across different trials, except for the $PV5$ case, shows minimal variance, and there is no clear distinction between cases where DNA or RNA viruses are excluded. This supports the hypothesis that proteins involved in chromatin structure may play a crucial role in distinguishing between the oncogenic and non-oncogenic cases.

Based on these considerations, it can be concluded that the classification model may not heavily rely on the type of RNA or DNA viruses. This provides support for the hypothesis that the model can identify oncogenic-related features beyond viral types.

### 10 GO enrichment analysis results, common nodes between all oncogenic layers

In the following we report the complete list of the 30 proteins obtained from the LVCs analysis described in sec. *Identifying the core of nodes biologically relevant for oncogenesis* from the article:

`RNF4, DAXX, PARP1, TP53, MDM2, CREBBP, ABL1, HMGA1, H2BC21, PIN1, FBXW7, PML, TRIM25, MAP3K1, MDM4,`
`UBE2I, NKX2-1, SMAD3, TP73, PPM1D, HNRNPL, VIRMA, SMAD2, CTBP1, CBX4, ACTBL2, RANBP9, SUMO2, SKI, PIAS1`

The results of the functional enrichment analysis based on Gene Ontology, encompassing Molecular Functions, Biological Processes and Cellular Components, are reported in Tab. 2.

### 11 GO enrichment analysis results, highly relevant proteins from perceptron weights analysis

In the following we report the complete list of the 81 proteins obtained from the machine learning classification task analysis described in sec. *Classification of oncogenic vs non-oncogenic virus-host interaction PPI networks using machine learning* from the article:

ASCC3, ATF1, ATP6AP2, ATRX, BST1, RTRAF, CBX3, CCNC, CEP170, CTBP1, CTNNA1, CTNND1, DCPS, DEFA5, DHX8,
DNMT3B, DNTTIP1, DYRK1B, E2F3, EAF1, EEF1A2, EMC2, SINHCAF, FTH1, GOPC, GPC6, MACROH2A1, H3F3A, HDAC8,
HES1, HIPK2, H2BC5, H3C1, H2AC18, H3C15, H3-7, H4C16, HSPG2, ID3, IDE, ITPR1, IWS1, PUM3, LGALS3BP, LRP2,
LYPD3, LZTS2, MAGED1, MECP2, MEF2D, MORF4L1, MTCH2, MYB, NCK2, NKX2-1, NLK, NPSR1, PDK1, PHF12, PKNOX1,
PLD1, PPM1D, RRM1, RSRC1, SAP30L, SENP1, SNX17, SP100, SSR1, SUV39H2, TAF6L, TCF12, TMED10, TNPO1,
TP53INP1, TRAF1, TRIM29, UBA2, WDR31, WNT3A, ZNF217

The results of the functional enrichment analysis based on Gene Ontology, encompassing Molecular Functions, Biological Processes and Cellular Components, are reported in Tab. 3.

### 12 GO enrichment analysis results, highly relevant proteins from perceptron weights analysis with different target set

In the following we report the complete list of the 184 proteins obtained from the machine learning classification task analysis described in sec. *Specificity on oncogenic non-oncogenic classification task.* from the article:

ACAD9, ADSL, AP1M1, ATP5ME, ATP5F1D, ATP5PF, ATP5MF, BST1, C15orf48, C4A, CCL7, CD1C, CD276, CD38,
COX4I1, COX6B1, CREB3, DEFA5, DHX37, DYNLT1, EXOSC6, F3, FCGR2B, FOXRED1, GDI1, GOLT1B, GUF1, H2BC5,
HLA-B, HLA-C, HLA-E, HLA-F, HLA-G, IDE, KRT1, KRT10, KRT12, KRT13, KRT14, KRT16, KRT19, KRT2, KRT20, KRT23,
KRT24, KRT25, KRT26, KRT27, KRT28, KRT3, KRT32, KRT33A, KRT33B, KRT34, KRT35, KRT36, KRT37, KRT38, KRT39,
KRT4, KRT5, KRT6A, KRT6B, KRT6C, KRT7, KRT71, KRT72, KRT73, KRT74, KRT75, KRT76, KRT77, KRT78, KRT79, KRT8,
KRT80, KRT81, KRT82, KRT83, KRT84, KRT85, KRT86, KRT9, LAP3, LTA, LTB, LTBR, LYPD3, MELTF, MRPL4, MSLN,
MT-CYB, MT-ND1, MT-ND2, MT-ND3, MT-ND4, MT-ND4L, MT-ND5, MT-ND6, NDUFA1, NDUFA10, NDUFA11, NDUFA12, NDUFA13,
NDUFA2, NDUFA3, NDUFA6, NDUFA7, NDUFA9, NDUFAF1, NDUFAF2, NDUFAF3, NDUFAF4, NDUFAF5, NDUFAF6, NDUFAF7,
NDUFB1, NDUFB10, NDUFB11, NDUFB2, NDUFB3, NDUFB4, NDUFB5, NDUFB6, NDUFB7, NDUFB8, NDUFB9, NDUFC1, NDUFS3,
NDUFS4, NDUFS5, NDUFS6, NDUFS8, NDUFV1, NDUFV2, NDUFV3, NOC4L, NOP14,

NPSR1, PECAM1, PGD, PLCB1, PLCB2,
PLCB3, PMPCB, RAB14, RAB9A, RAD51C, REN, RPL29, RPL36AL, RPP38, SELL,
SFXN5, SLC25A28, TAMM41, TARS1, TECR,
TFCP2, THBD, TIMMDC1, TMEM126B, TNFRSF11A, TNFRSF12A, TNFRSF13C,
TNFRSF4, TNFRSF9, TNFSF12, TNFSF14, TRIO,
TSR1, TXNRD1, UCHL5, UQCR10, UQCR11, UQCRB, UQCRFS1, UQCRH, UQCRQ,
USP43, WDR3, WWP2, YBEY

The results of the functional enrichment analysis based on Gene Ontology, encompassing Molecular Functions, Biological Processes and Cellular Components, are reported in Tab. 4.

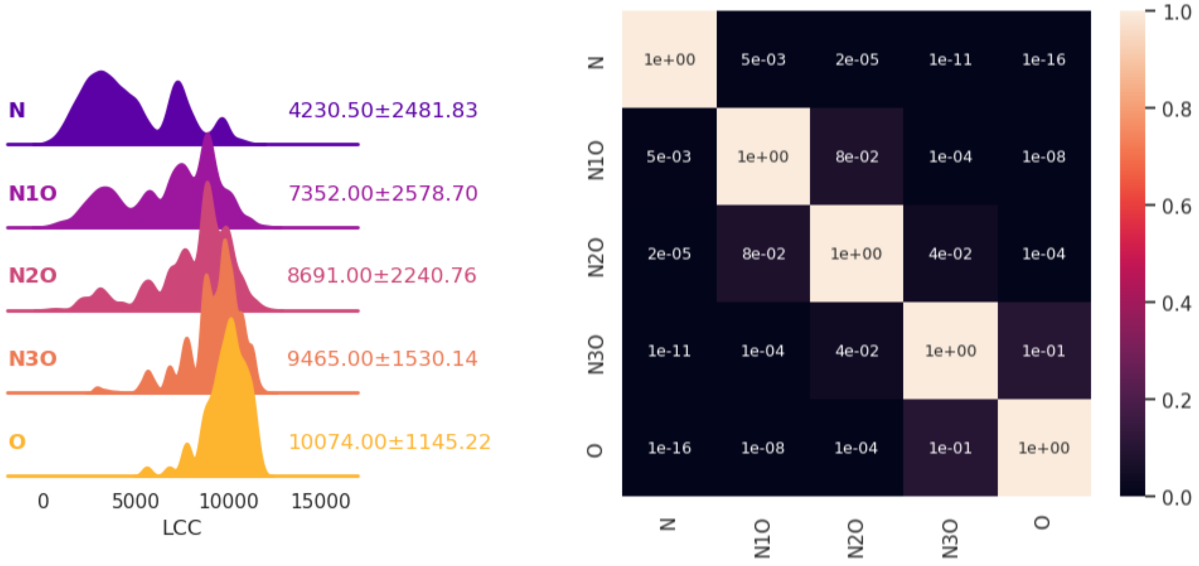


Figure 1: On the left the normalized distribution plots depict the LCC size for samples belonging to different combination sets. The median and standard deviation values are provided for each distribution, offering insights into the central tendency and variability of the respective variables. On the left the heatmap displays the p-values obtained from the Wilcoxon-Ranksum test applied to all possible pairs of distributions.

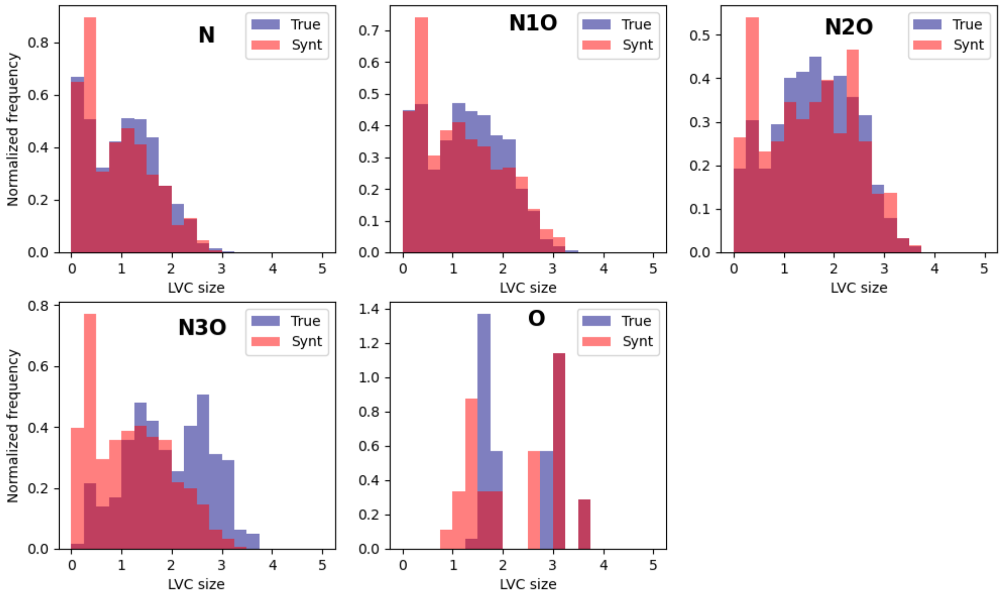


Figure 2: Each figure represent the distributions of the LVC sizes. The blue histogram is associated to a set of samples extracted from the combination set build with the original data reported bold in the figures, while the red one takes into account samples from the corresponding combination sets created from 500 realization of the randomized model.

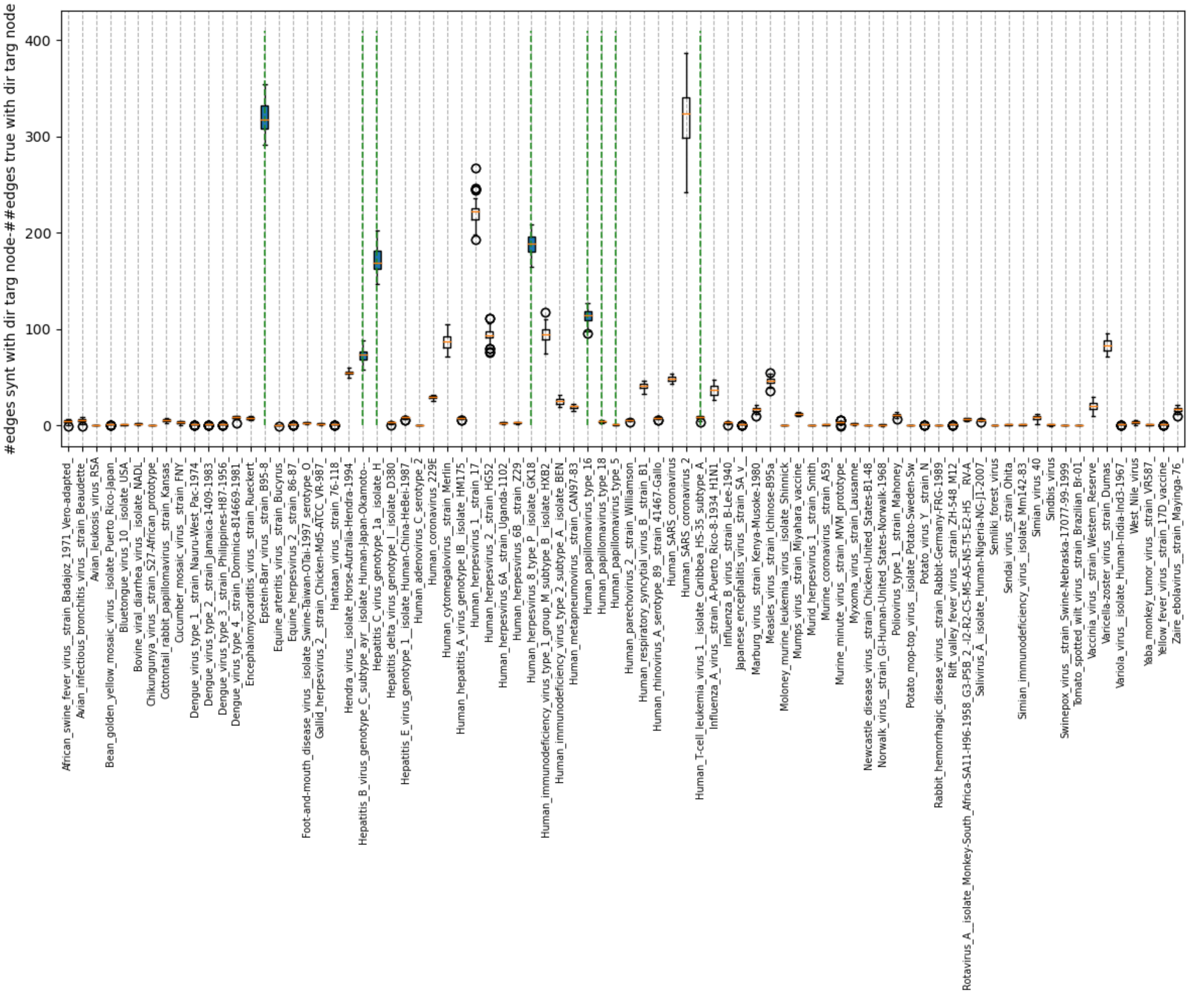


Figure 3: Difference between the number of edges connecting at least one node directly targeted by a virus in the corresponding virus-host interaction network extracted from a randomization of the original human PPI network and the number of edges connecting at least one node directly targeted by a virus in the corresponding virus-host interaction network extracted from the original human PPI network

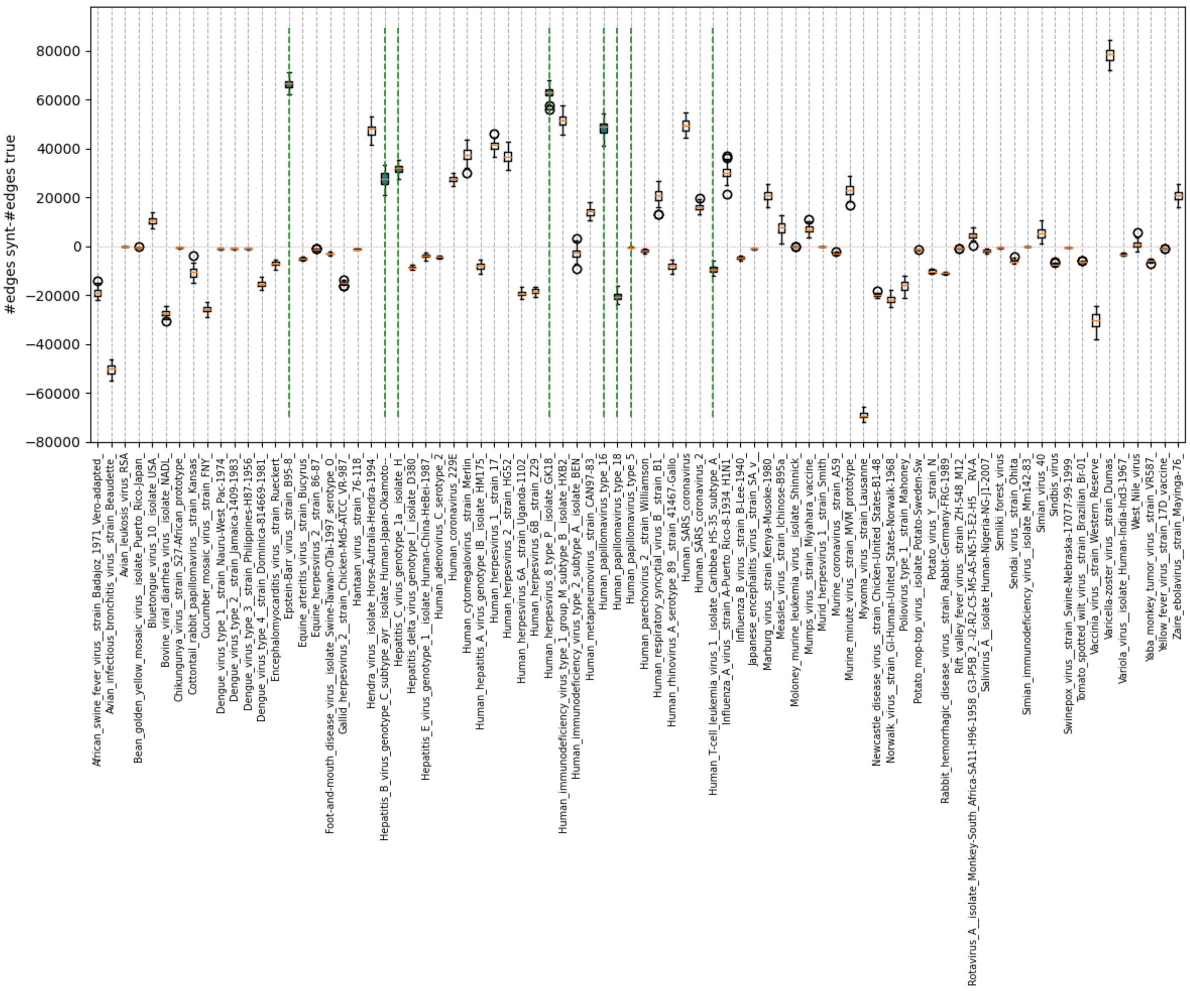


Figure 4: Difference between the total number of edges in the corresponding virus-host interaction network extracted from a randomization of the original human PPI network and the number of edges in the corresponding virus-host interaction network extracted the original human PPI network

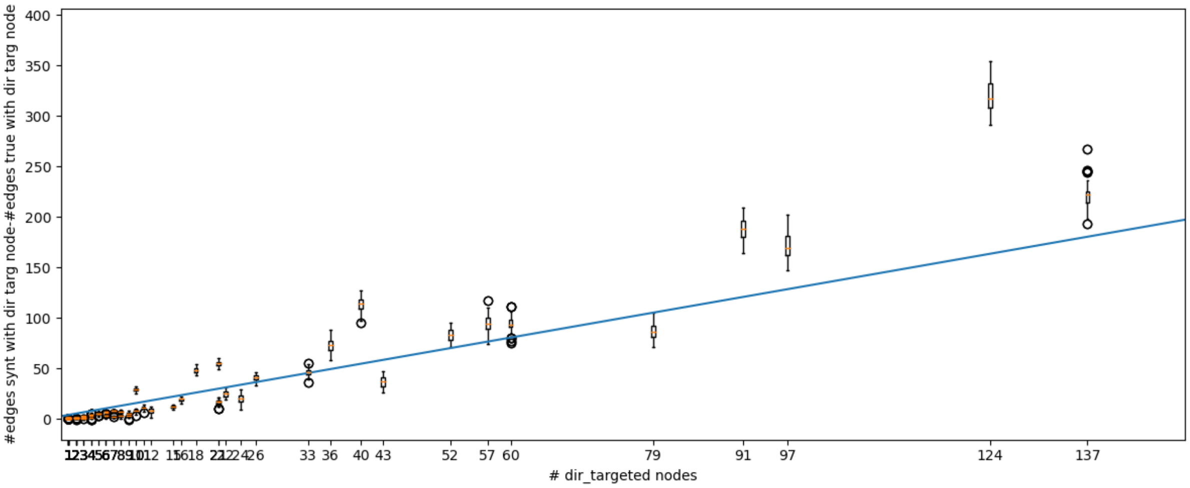


Figure 5: Difference between the number of edges connecting at least one node directly targeted by a virus in the corresponding virus-host interaction network extracted from a randomization of the original human PPI network and the number of edges connecting at least one node directly targeted by a virus in the corresponding virus-host interaction network extracted from the original human PPI network vs number of directly targeted human proteins by the virus

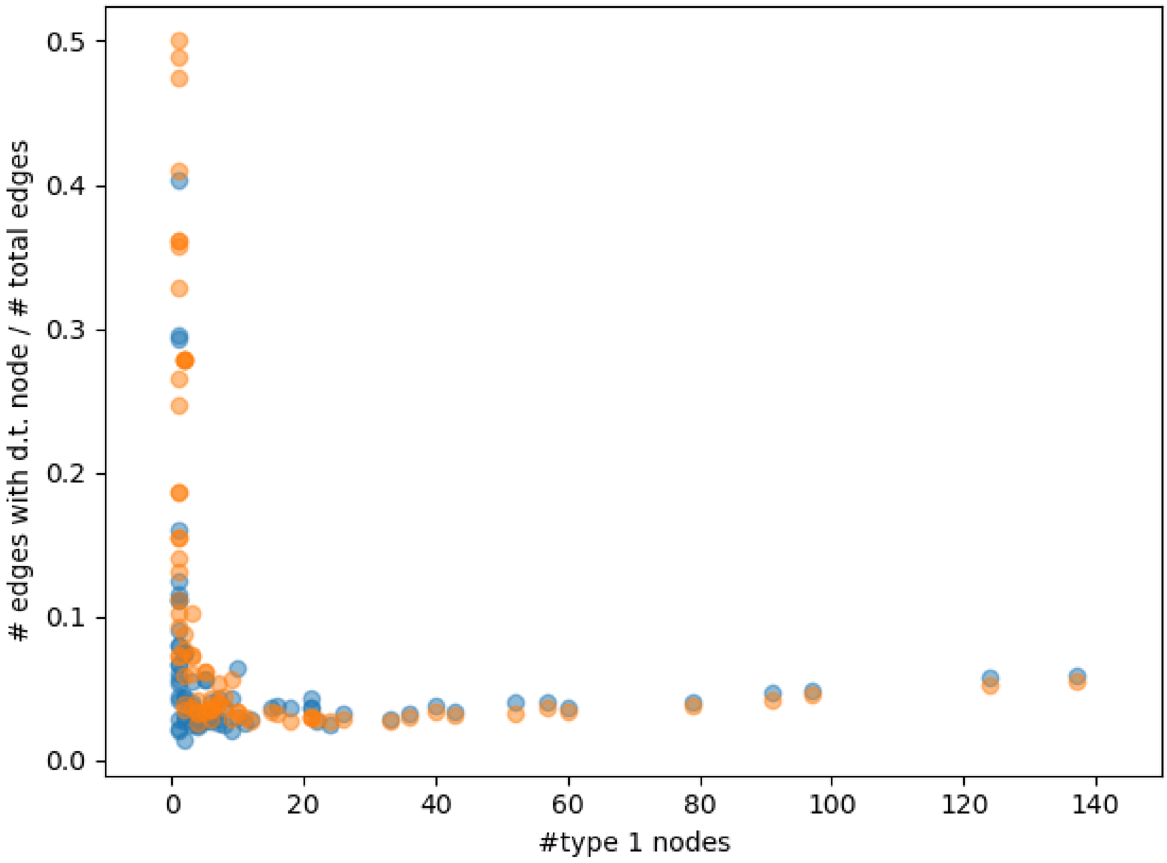


Figure 6: Ratio between number of edges containing a directly targeted node by a virus and the total number of edges of the virus-host interaction PPI network, as a function of the number of directly targeted nodes by the virus.

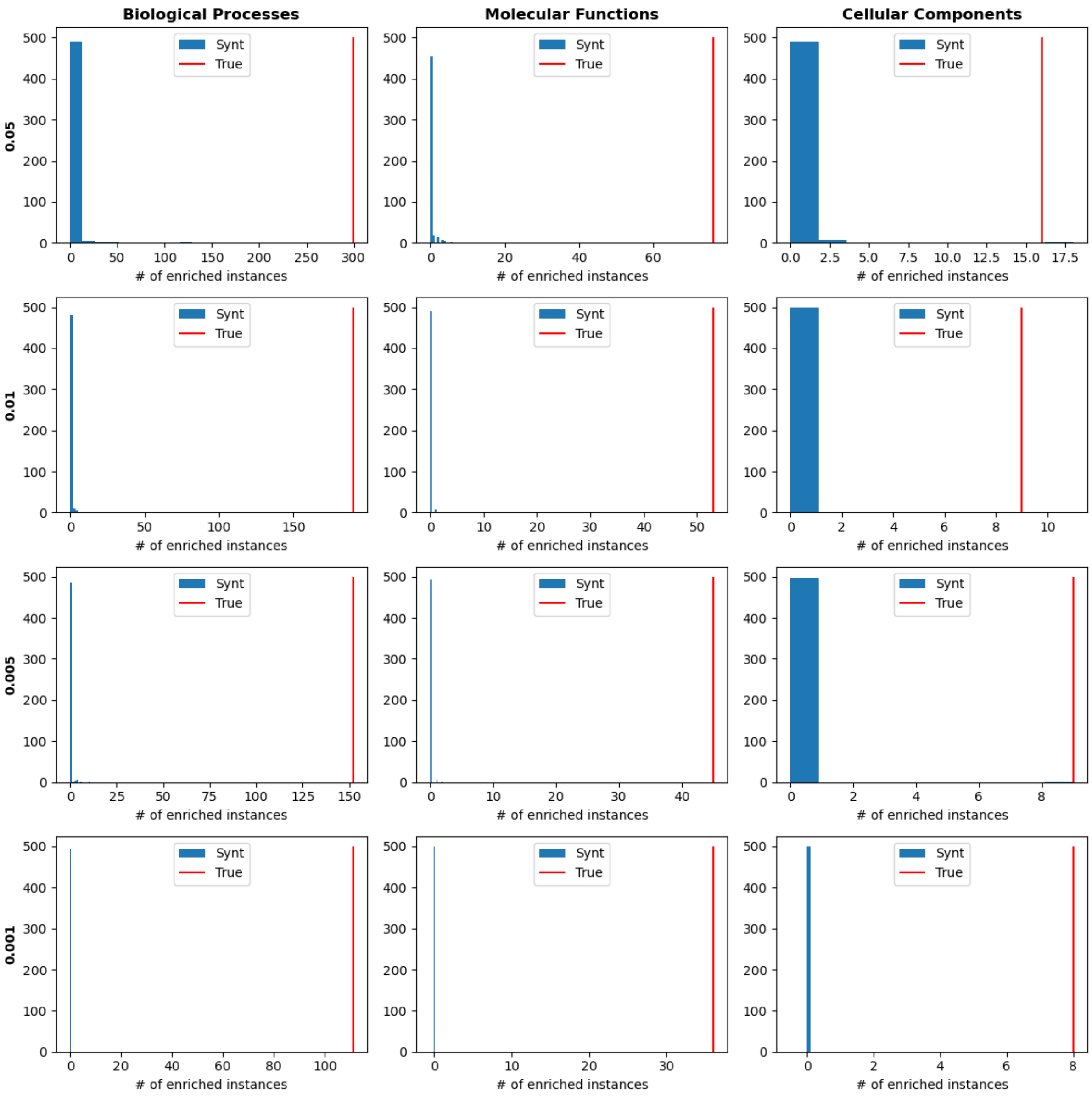


Figure 7: The figures reports the number of enriched GO instances by the set of proteins composing the LVCs of the multilayer network composed by 8 layers corresponding to the 8 PPINs corresponding to the oncogenic viruses. The blue histograms describes the distributions of the results obtained starting from 500 realization of the randomized model, while the red line indicated the value extracted from the original dataset. Each column represent a different GO category over which the enrichment analysis is performed, while the rows indicated the FDR corrected p-value threshold under which an instance is considered to be enriched.

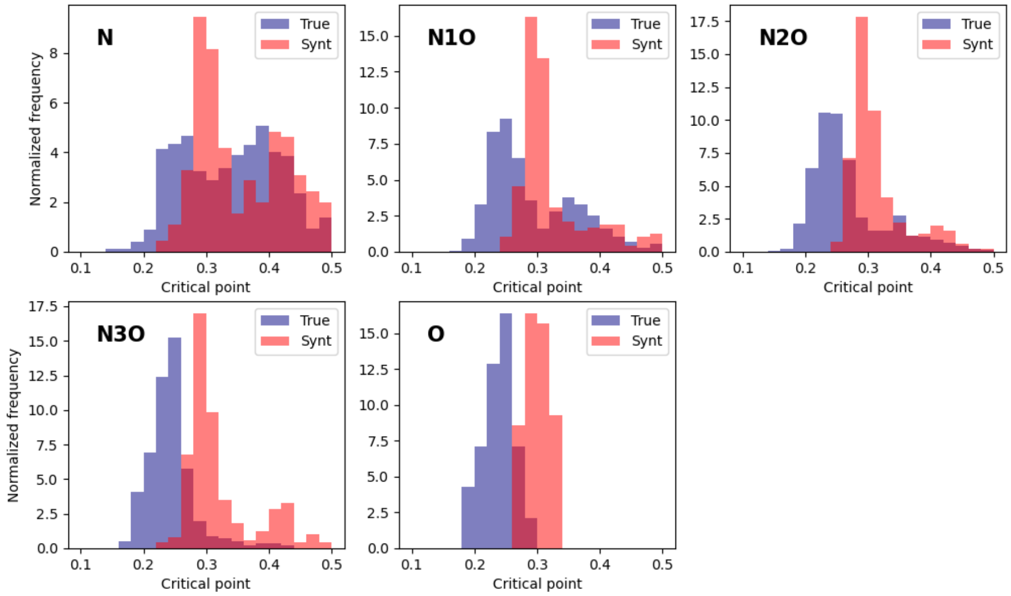


Figure 8: Each figure represent the distributions of the critical point of the multilayer page-rank ordered node percolation process. The blue histogram is associated to a set of samples extracted from the combination set build with the original data reported bold in the figures, while the red one takes into account samples from the corresponding combination sets created from 500 realization of the randomized model.

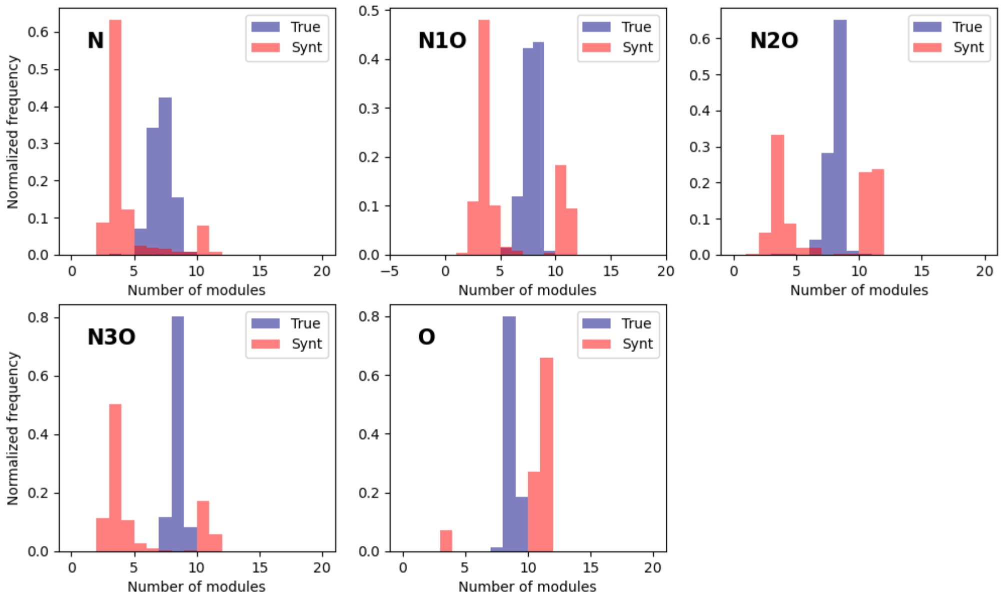


Figure 9: Each figure represent the distributions of the number of modules of the multilayer networks computed by using the DCSBM algorithm. The blue histogram is associated to a set of samples extracted from the combination set build with the original data reported bold in the figures, while the red one takes into account samples from the corresponding combination sets created from 500 realization of the randomized model.

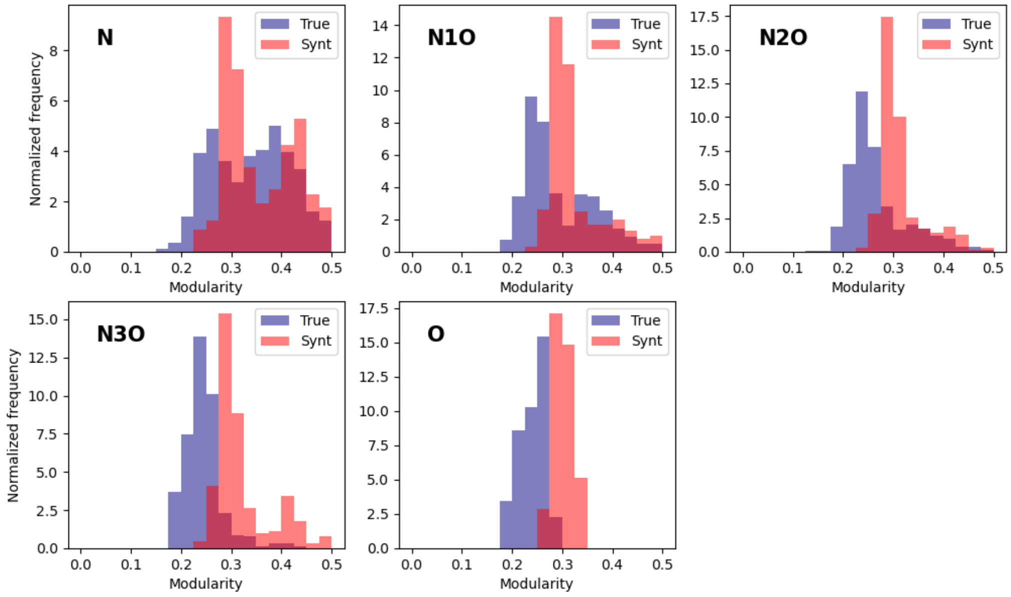


Figure 10: Each figure represent the distributions of the modularity of the multilayer networks computed by using the DCSBM algorithm. The blue histogram is associated to a set of samples extracted from the combination set build with the original data reported bold in the figures, while the red one takes into account samples from the corresponding combination sets created from 500 realization of the randomized model.

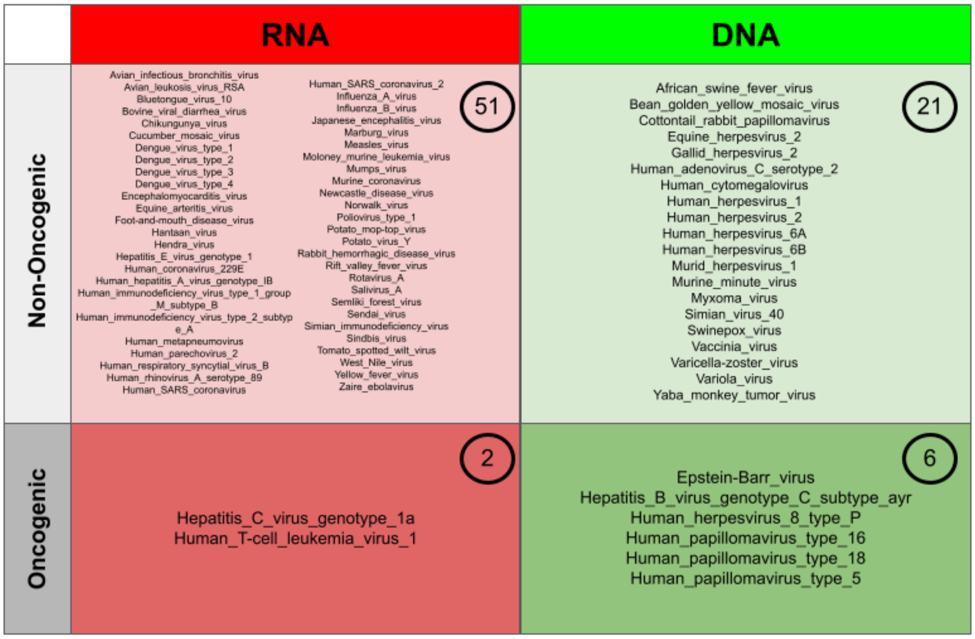


Figure 11: Classification of the viruses in the dataset based on their genetic material (DNA vs RNA) and the oncogenic feature. In the top right of each box is reported the total number of viruses with that specific set of features

| Trial Name | Excluded Onco Virus | Train Acc | Val Acc | OncoTest pred |
|---|---|---|---|---|
| EB | Epstein-Barr | 0.923 | 0.931 | 0.962 |
| HBC | Hepatitis B gen. C, ayr | 0.916 | 0.9204 | 0.670 |
| HC1 | Hepatitis C gen. 1a | 0.931 | 0.920 | 0.886 |
| HV8P | Hum. herpesvirus 8 type P | 0.923 | 0.898 | 0.969 |
| PV16 | Hum. papillomavirus type 16 | 0.918 | 0.911 | 0.736 |
| PV18 | Hum. papillomavirus type 18 | 0.917 | 0.916 | 0.833 |
| PV5 | Hum. papillomavirus type 5 | 0.932 | 0.932 | 0.402 |
| TL1 | Hum. T-cell leukemia 1 | 0.915 | 0.920 | 0.860 |

Table 1: The table presents the performance results of individual Perceptron models trained using datasets in which samples containing specific oncogenic viruses were excluded, specifically the training and validation accuracy at the end of each model training. The *OncoTest pred* column contains the accuracy values of the predictions performed over the samples containing the excluded oncogenic virus PPI layer.

| Name | p-value | Bonferroni | FDR B&H | FDR B&Y |
|---|---|---|---|---|
| **MOLECULAR FUNCTIONS** | | | | |
| DNA-binding transcription factor binding | 1.165E-15 | 3.087E-13 | 3.087E-13 | 1.901E-12 |
| transcription factor binding | 4.734E-14 | 1.254E-11 | 6.272E-12 | 3.863E-11 |
| SUMO transferase activity | 8.266E-14 | 2.190E-11 | 7.301E-12 | 4.497E-11 |
| RNA polymerase II-specific DNA-binding transcription factor binding | 5.525E-13 | 1.464E-10 | 3.660E-11 | 2.254E-10 |
| ubiquitin protein ligase binding | 1.293E-12 | 3.426E-10 | 6.852E-11 | 4.220E-10 |
| ubiquitin-like protein ligase binding | 2.472E-12 | 6.552E-10 | 1.092E-10 | 6.725E-10 |
| **BIOLOGICAL PROCESS** | | | | |
| negative regulation of DNA-templated transcription | 9.778E-14 | 2.282E-10 | 1.013E-10 | 8.438E-10 |
| negative regulation of RNA biosynthetic process | 1.188E-13 | 2.774E-10 | 1.013E-10 | 8.438E-10 |
| negative regulation of nucleobase-containing compound metabolic process | 1.302E-13 | 3.038E-10 | 1.013E-10 | 8.438E-10 |
| positive regulation of DNA-templated transcription | 3.789E-13 | 8.844E-10 | 1.872E-10 | 1.560E-9 |
| positive regulation of RNA biosynthetic process | 4.011E-13 | 9.361E-10 | 1.872E-10 | 1.560E-9 |
| negative regulation of RNA metabolic process | 5.738E-13 | 1.339E-9 | 1.934E-10 | 1.612E-9 |
| **CELLULAR COMPONENTS** | | | | |
| nuclear body | 4.860E-16 | 6.902E-14 | 6.902E-14 | 3.821E-13 |
| PML body | 7.477E-14 | 1.062E-11 | 5.308E-12 | 2.939E-11 |
| transcription regulator complex | 1.688E-11 | 2.397E-9 | 7.989E-10 | 4.423E-9 |
| chromatin | 3.287E-8 | 4.668E-6 | 1.167E-6 | 6.461E-6 |
| protein-DNA complex | 5.967E-8 | 8.473E-6 | 1.695E-6 | 9.382E-6 |
| transcription repressor complex | 1.373E-7 | 1.950E-5 | 3.250E-6 | 1.799E-5 |

Table 2: Results of the GO functional enrichment analysis performed over the set of proteins composing the LVC of the multilayer network composed the 8 PPINs associated to the oncogenic viruses. The analysis was performed using ToppGene

| Name | p-value | Bonferroni | FDR B&H | FDR B&Y |
| --- | --- | --- | --- | --- |
| **MOLECULAR FUNCTIONS** | | | | |
| protein dimerization activity | 3.139E-9 | 1.224E-6 | 1.224E-6 | 8.012E-6 |
| structural constituent of chromatin | 1.151E-8 | 4.488E-6 | 2.244E-6 | 1.469E-5 |
| protein heterodimerization activity | 2.283E-8 | 8.903E-6 | 2.968E-6 | 1.942E-5 |
| transcription factor binding | 6.390E-8 | 2.492E-5 | 6.230E-6 | 4.078E-5 |
| protein domain specific binding | 8.869E-8 | 3.459E-5 | 6.918E-6 | 4.528E-5 |
| chromatin binding | 2.013E-7 | 7.850E-5 | 1.308E-5 | 8.562E-5 |
| **BIOLOGICAL PROCESS** | | | | |
| negative regulation of transcription by RNA polymerase II | 1.173E-9 | 3.100E-6 | 2.883E-6 | 2.438E-5 |
| negative regulation of gene expression, epigenetic | 2.182E-9 | 5.766E-6 | 2.883E-6 | 2.438E-5 |
| peptidyl-amino acid modification | 6.448E-9 | 1.704E-5 | 4.366E-6 | 3.692E-5 |
| chromatin remodeling | 7.127E-9 | 1.884E-5 | 4.366E-6 | 3.692E-5 |
| heterochromatin formation | 8.259E-9 | 2.183E-5 | 4.366E-6 | 3.692E-5 |
| heterochromatin organization | 1.664E-8 | 4.399E-5 | 7.332E-6 | 6.201E-5 |
| **CELLULAR COMPONENTS** | | | | |
| chromatin | 2.545E-12 | 8.628E-10 | 8.628E-10 | 5.526E-9 |
| protein-DNA complex | 8.203E-12 | 2.781E-9 | 1.390E-9 | 8.906E-9 |
| histone deacetylase complex | 1.343E-9 | 4.554E-7 | 1.518E-7 | 9.723E-7 |
| nucleosome | 4.091E-9 | 1.387E-6 | 3.467E-7 | 2.221E-6 |
| nuclear protein-containing complex | 2.215E-7 | 7.509E-5 | 1.502E-5 | 9.618E-5 |
| nuclear chromosome | 7.149E-7 | 2.424E-4 | 4.039E-5 | 2.587E-4 |

Table 3: Results of the GO functional enrichment analysis performed over the set of highly relevant proteins from the oncogenic/non-oncogenic machine learning classification procedure. The analysis was performed using ToppGene

| Name | p-value | Bonferroni | FDR B&H | FDR B&Y |
|---|---|---|---|---|
| **MOLECULAR FUNCTIONS** | | | | |
| oxidoreduction-driven active transmembrane transporter activity | 1.363E-65 | 5.288E-63 | 5.288E-63 | 3.458E-62 |
| NADH dehydrogenase (ubiquinone) activity | 1.101E-64 | 4.271E-62 | 2.136E-62 | 1.397E-61 |
| NADH dehydrogenase (quinone) activity | 4.918E-64 | 1.908E-61 | 6.360E-62 | 4.159E-61 |
| NAD(P)H dehydrogenase (quinone) activity | 2.910E-62 | 1.129E-59 | 2.823E-60 | 1.846E-59 |
| NADH dehydrogenase activity | 1.011E-61 | 3.923E-59 | 7.846E-60 | 5.131E-59 |
| oxidoreductase activity, acting on NAD(P)H, quinone or similar compound as acceptor | 3.810E-56 | 1.478E-53 | 2.464E-54 | 1.611E-53 |
| **BIOLOGICAL PROCESS** | | | | |
| intermediate filament organization | 7.791E-84 | 1.869E-80 | 1.869E-80 | 1.563E-79 |
| intermediate filament cytoskeleton organization | 2.319E-75 | 5.563E-72 | 2.781E-72 | 2.325E-71 |
| intermediate filament-based process | 4.475E-75 | 1.073E-71 | 3.578E-72 | 2.991E-71 |
| mitochondrial respiratory chain complex I assembly | 1.673E-68 | 4.014E-65 | 8.028E-66 | 6.712E-65 |
| NADH dehydrogenase complex assembly | 1.673E-68 | 4.014E-65 | 8.028E-66 | 6.712E-65 |
| proton motive force-driven mitochondrial ATP synthesis | 2.456E-67 | 5.892E-64 | 9.820E-65 | 8.209E-64 |
| **CELLULAR COMPONENTS** | | | | |
| respiratory chain complex | 8.125E-77 | 2.494E-74 | 2.494E-74 | 1.573E-73 |
| mitochondrial respirasome | 6.378E-76 | 1.958E-73 | 9.791E-74 | 6.174E-73 |
| respirasome | 1.875E-73 | 5.755E-71 | 1.918E-71 | 1.210E-70 |
| mitochondrial respiratory chain complex I | 5.300E-71 | 1.627E-68 | 2.712E-69 | 1.710E-68 |
| NADH dehydrogenase complex | 5.300E-71 | 1.627E-68 | 2.712E-69 | 1.710E-68 |
| respiratory chain complex I | 5.300E-71 | 1.627E-68 | 2.712E-69 | 1.710E-68 |

Table 4: Results of the GO functional enrichment analysis performed over the set of highly relevant proteins from the machine learning classification procedure in which the set of target virus was changed. The analysis was performed using ToppGene

### References and Notes


[1] De Domenico, M. *et al.* Mathematical formulation of multilayer networks. *Phys. Rev. X* **3**, 041022 (2013). URL `https://link.aps.org/doi/10.1103/PhysRevX.3.041022`.

[2] De Domenico, M., Solé-Ribalta, A., Omodei, E., Gómez, S. & Arenas, A. Ranking in interconnected multilayer networks reveals versatile nodes. *Nature Communications* **6** (2015). URL `http://dx.doi.org/10.1038/ncomms7868`.

[3] De Domenico, M., Porter, M. A. & Arenas, A. Muxviz: a tool for multilayer analysis and visualization of networks. *Journal of Complex Networks* **3**, 159–176 (2014). URL `http://dx.doi.org/10.1093/comnet/cnu038`.
